# G-quadruplex ligand RHPS4 compromises cellular radio-resistance by blocking the increase in mitochondrial mass and activity induced by ionising irradiation

**DOI:** 10.1101/2024.10.24.619985

**Authors:** S. Tricot, C. Siberchicot, I. Bontemps, C. Desmaze, G. Kratassiouk, M. Vandamme, G. Pinna, J.Pablo Radicella, G. Lenaers, J. Lebeau, A. Campalans

## Abstract

A major challenge in radiotherapy is to enhance tumor cell sensitivity to radiation while minimizing damage to healthy tissues. Ionising radiation (IR) induces mitochondrial DNA (mtDNA) alterations that can impair mitochondrial function and cell survival. Since mitochondria play a key role in tumor cell proliferation, they represent a promising therapeutic target for cancer treatment. In this study, we characterized the impact of different IR sources on mitochondrial function in radioresistant cancer cells. Our findings revealed several adaptive responses that may contribute to radioresistance, including increased mtDNA content, mitochondrial mass, enhanced activity, and hyperfusion of the mitochondrial network. Notably, the use of mitochondrial-targeted G-quadruplex (G4) ligands, which block mtDNA replication and transcription, disrupted these responses, reducing cancer cell survival in an mtDNA-dependent manner. These results demonstrate that mitochondrial adaptations contribute to radioresistance and highlight mitochondria as a novel target for the radiosensitizing effects of G4-ligands, extending their potential beyond telomere destabilization.

## INTRODUCTION

Radiotherapy (RT), together with surgery and chemotherapy, is one of the major forms of cancer therapy. Its primary goal is to target malignant cells by inducing non-reparable DNA damage, while sparing the healthy tissue surrounding the tumor as much as possible. Despite the efficiency of radiotherapy in cancer treatment, the adaptive response of cancer cells to radiation-induced damage is a major problem being at the origin of therapeutic failure and relapse. Therefore, an intense research is dedicated to improve the therapeutic index of radiotherapy either by developing new irradiation strategies allowing a more precise targeting of the tumor (proton beam therapy, FLASH-RT) or by developing targeted drugs enhancing cancer cell radiosensitivity (Begg et al., 2011).

Although the effects of ionising radiation on nuclear DNA have been extensively studied, much less is known concerning its impact on mitochondrial DNA and function. Human cells contain multiple copies of mtDNA, a small circular molecule of 16 Kb that encodes 13 subunits of the mitochondrial respiratory chain together with rRNAs and tRNAs involved in their translation. mtDNA is therefore required to sustain oxidative phosphorylation and mitochondrial functions, essential for the proliferation of cancer cells (Wallace, 2012; Liu et al., 2023; Zaffaroni et al., 2022). Irradiation has been previously reported to increase the induction of the common deletion in mtDNA (Chen et al., 2018; Schilling-Toth et al., 2011; Prithivirajsingh et al., 2004) representing a loss of roughly 1/3 of the mtDNA molecule (4977 bp), containing 5 tRNA genes and 7 genes coding for subunits of the respiratory complexes (Yusoff et al., 2019). Fragmentation of the mitochondrial network has also been observed after irradiation, suggesting the induction of mitochondrial dysfunction a few hours after treatment (Jin et al., 2018). Furthermore, mitochondrial adaptation plays a crucial role in the development of chemo and radioresistance, hindering cancer treatment by potentially leading to tumor relapse and metastatsis (McCann et al., 2021).

Therefore, inducing mitochondrial dysfunction is emerging as a promising strategy in cancer treatment and many molecules have been developed targeting mitochondrial metabolism, mitochondrial redox signaling pathways, mitochondrial dynamics and mitochondrial DNA (mtDNA) (Dong et al., 2020). Several compounds have been identified to affect the stability and activity of mtDNA and it has been shown that elimination of mtDNA results in an inhibition of cancer cell proliferation therefore limiting tumorigenesis (Tan et al., 2015). An example are the recently developed small-molecule inhibitors of the human mitochondrial RNA polymerase POLRMT, that induce a defect in the biogenesis of the oxidative phosphorylation system resulting in the specific inhibition of cancer cell proliferation and tumor growth in a preclinical mouse model (Bonekamp et al., 2020).

One of the features of mtDNA is its high GC content and its GC strand asymmetry with a two-fold enrichment of guanines on one strand contributing to the very high density of sequences with the potential to form G-quadruplexes (G4). G4s are secondary structures formed in guanine rich regions by stacked sets of four guanines that have been involved in regulation of replication, transcription and telomere maintenance of nuclear DNA. Evidence indicate that G4 may also play important roles in the regulation of mitochondrial DNA replication and transcription (Sahayasheela et al., 2023; Wanrooij et al., 2010 and 2012) and being particularly enriched at breakpoints of pathogenic mtDNA deletions they are at the origin of mtDNA instability (Falabella et al., 2019; Doimo et al., 2023; Bharti et al., 2014; Dahal et al., 2021).

G4 structures have been identified as promising anti-cancer targets and several hundreds of G4 ligands with the capacity to recognize and stabilize these structures have been developed. G4 ligands can target multiple genomic regions and affect cancer cell proliferation by inducing telomere instability or stabilizing G4 at promoters therefore altering oncogene expression (Figueiredo et al., 2024). Radiosensitizing effects of G4 ligands leading to cell cycle arrest and apoptosis induction have been reported and explained by an increase in telomere dysfunction (Merle et al., 2015; Berardinelli et al., 2015). A very promising molecule is the G4 ligand RHPS4 that combined with irradiation reduces tumor mass and avoids tumor relapse by inducing telomere dysfunction, replicative stress and accumulation of nuclear DNA damage depending on the cellular context (Berardinelli et al., 2019). Interestingly, recent findings have shown that RHPS4 can also localize to mitochondria and affect mitochondrial transcription and replication resulting in a loss of mtDNA and a depletion of respiratory complexes (Falabella et al., 2019; Fallabella et al., 2020; Doimo et al., 2023).

In this study we have characterized the effects of ionising radiation on the mitochondrial network and activity and shown an adaptive response resulting in an increase in mtDNA copy number, mitochondrial mass and activity together with the hyperfusion of mitochondrial network, potentially contributing to cellular radioresistance. The mitochondrially localized G4 ligand RHPS4 compromises the mitochondrial adaptation induced by irradiation and increases cellular radiosensitivity in a mtDNA dependent manner. Our results not only bring further insights on the effects of radiation on mitochondria and the molecular mechanisms involving G4 in mtDNA but also highlight a novel therapeutical application of this promising small molecule in cancer treatment.

## RESULTS

### Increase in mitochondrial mass and activity induced by ionising radiation

In order to evaluate the impact of ionising radiation on mitochondrial content, mtDNA copy number was monitored by qPCR at different times after a single dose of gamma irradiation at 6 Gy. In order to evaluate the eventual accumulation of molecules with the common deletion in irradiated cells we selected probes against the mitochondrial genes ND1, COX2 and COX3 and two intergenic regions located throughout the mtDNA molecule, inside or outside the common deletion. An increase in the levels of mtDNA was clearly observed 96H after a single irradiation at 6 Gy in surviving cells for all the probes used, indicating an amplification of the entire mtDNA molecule (Figure 1A).

**Figure 1.**
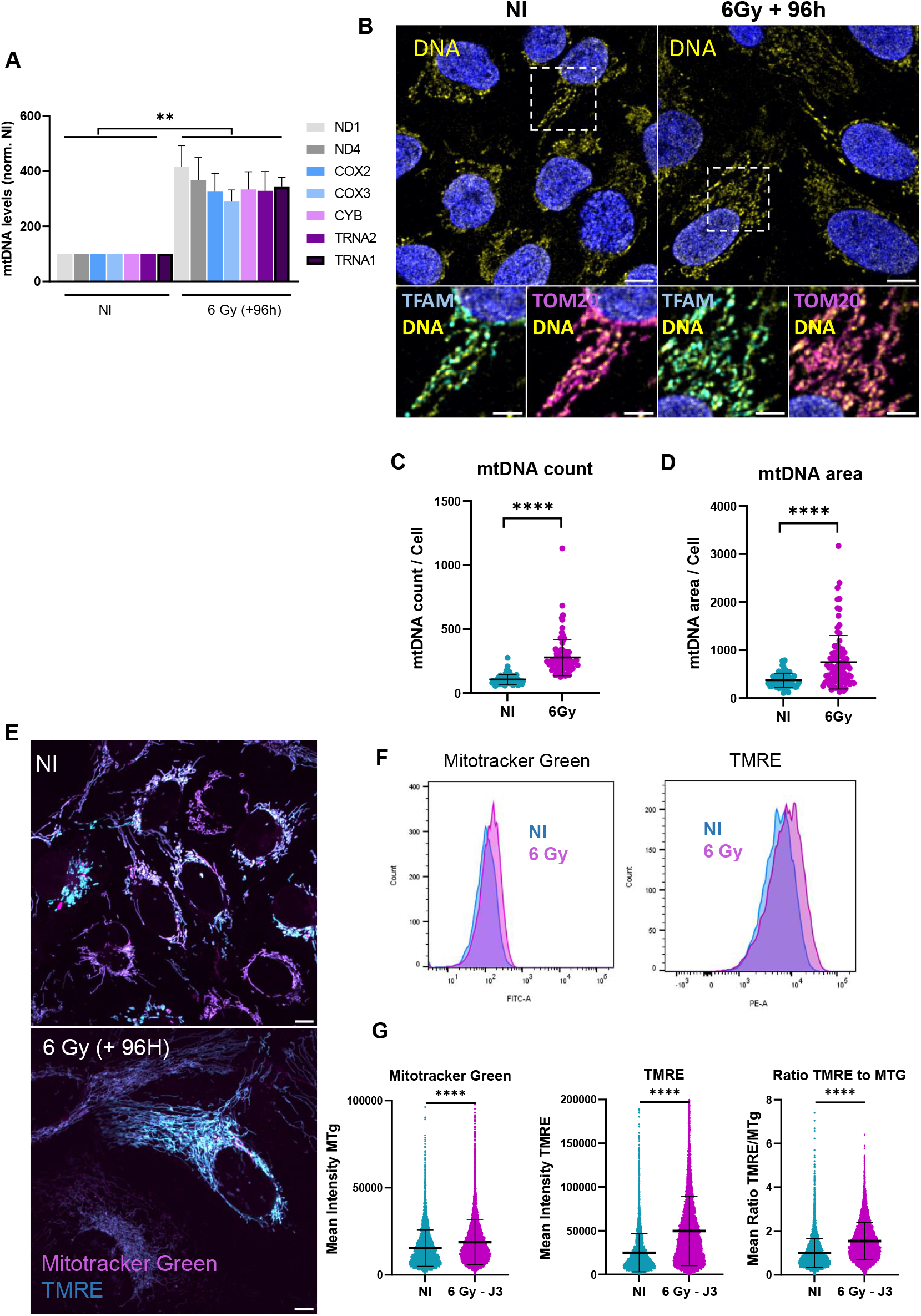
Increase in mitochondrial mass and activity induced by Ionising radiation. U2OS cells were irradiated by gamma irradiation (6Gy) and compared with no irradiated cells. **A)** mitochondrial genes ND1, ND4, COX2, COX3, CYB, TRNA1 and TRNA2 were quantified by qPCR 96 h after irradiation. Statistical analysis was performed for five independent experiments with one-way ANOVA (**<0.005). **A)**Confocal microscopy images of U2OS cells stained with anti-DNA (yellow), anti-TFAM (cyan) and TOM20 (magenta). Nuclear DNA was stained with DAPI (blue). Scale bar is 10µm and 5µm for insets. **C)**number of nucleoids and **D)** the area of mtDNA per cell was quantified from more than 1000 cells from 4 independent experiments. **E)** Confocal microscopy images of U2OS living cells 96h after irradiation. Mitochondria were stained with MitoTracker Green (magenta) and TMRE (cyan), the scale bar is 10µm. **F)** TMRE and MitoTracker Green probes were used to measure mitochondrial membrane protential and mitochondrial mass, respectively in U2OS cell by cytometry 3 days after irradiation (6Gy). **G)** Representative quantification of cytometry results for single cells MitoTracker Green, TMRE and the ratio TMRE to MitoTracker Green (MTG). 10 000 cells were analysed for each condition. One representative experiment out of 3 is shown. Statistical analysis were performed with GraphPad using Mann-Whitney test (***P ≤ 0.001).

Mitochondrial nucleoids were visualized by immunofluorescence using an antibody against DNA, unveiling a signal specifically located within the mitochondrial network labeled with MitoTracker Red, and colocalizing with TFAM, a core protein of the mitochondrial nucleoid, in both irradiated and non-irradiated cells (Figure 1B). In agreement with the results from qPCR, an increase in the number and the surface of nucleoids per cell was observed in irradiated cells (Figure 1C-D).

The increase in mtDNA correlated with an increase in mitochondrial mass and membrane potential measured using MitoTracker Green and TMRE probes respectively (Figure 1E-G). Microscopy images confirm the specificity of the mitochondrial staining (Figure 1E). Even if both signals, MitoTracker Green and TMRE, increased in irradiated cells, the increase was more pronounced for TMRE, resulting in higher ratios TMRE / MitoTracker Green and thus indicating a higher mitochondrial activity in irradiated cells (Figure 1F-G).

Both microscopy and cytometry analyses revealed a gradual increase in cell size induced by ionising radiation (Figure S1A), with a proportion of large cells reaching up to 60% of the population four days after irradiation. Interestingly, the irradiation induced increase in mitochondrial mass and activity (Figure S1C), as well as the mtDNA copy number (Figure S6E) was specifically observed in these larger cells. Therefore, the increase in the proportion of large cells likely accounts for the gradual rise in mtDNA levels observed in the days following irradiation (Figure S1B).

### Ionising radiation induces hyperfusion of mitochondrial network

Mitochondrial dynamics involve the continuous processes of fusion and fission, which are crucial for maintaining mitochondrial function and cellular homeostasis. While fusion allows merging of mitochondrial contents and thus compensates for damaged components, fission allows mitochondrial division and removal of damaged mitochondria by mitophagy. The balance between fusion and fission ensures proper mitochondrial size, number, and function, which are essential for cellular energy production and overall cell viability (Tilokani et al., 2018; Youle et al., 2012).

Quantitative analysis of mitochondrial morphology in both living and fixed cells revealed an increase in mitochondrial footprint and in the number and the surface of branches (contacts between different parts of the network) induced by irradiation, clearly indicating an hyperfusion of mitochondrial network (Figure 2A-C; Figure S2). Mitochondrial hyperfusion has been previously described as an adaptation phenomenon allowing cells to survive to different kind of stresses such as nutrient deprivation or oxidative stress (Tondera et al., 2009). This adaptive response helps maintaining cellular energy balance and protects cells against apoptosis and therefore could contribute to cellular radioresistance.

**Figure 2.**
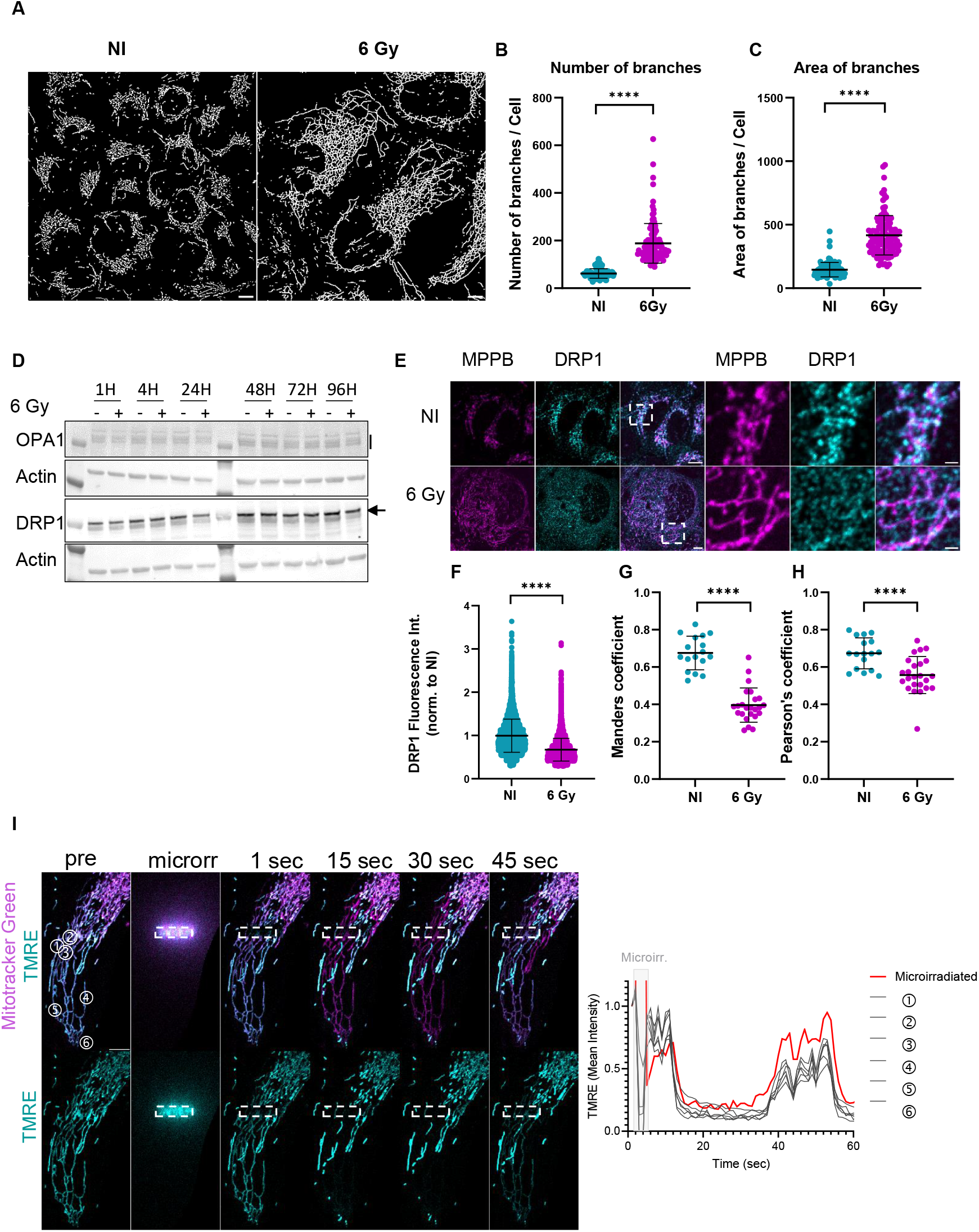
Ionising radiation induces functional hyperfusion of mitochondrial network. U2OS cells were gamma irradiated (6Gy) and compared with no irradiated cells. **A)** Mitochondria were stained with MitoTracker Red and Binary images were generated and analysed by applying the Kernel algorithm (Koopman et al., 2005). **B)** Number of mitochondrial branches and **C)** Area of branches per cell were analyse by ImageJ, see material and methods and in Fig.S2 for further details. More than 100 cells were analysed from 4 independent experiments. **D)** Levels of OPA1 and DRP1 were analysed by Western blot the indicated times after irradiation. Actin was used as a loading marker. **E)** Confocal microscopy images of U2OS cells stained with DRP1 (cyan) and PMPCB (magenta). Scale bar is 10µm and 2µm for insets. **F)** DRP1 fluorescence intensity in PMPCB mitochondrial mask was analysed and normalized to non-irradiated cells. More than 50 cells were analysed for each condition. Results from one representative experiment out of three, **G)** Mander’s **H)** Pearson correlation factors between DRP1 and PMPCB were determined with the JACOP plugin (imageJ). 17 NI and 24 Irradiated cells were analysed from three independent experiments. **I)** Microirradiation was performed with a 561 nm laser for 50 msec on the region indicated by the dotted square in cells stained with MitoTracker Green and TMRE. Cells were imaged at 1 image/sec during 2 min. The graph shows the quantification of the TMRE signal in the MitoTracker mask for mitochondria at the irradiated region (red) and areas located several microns away (black). A representative cell is shown. Scale bar 10 µm. Statistical analysis was performed in GraphPad using Mann-Whitney test (***P ≤ 0.001).

While no difference was observed after irradiation in the global amounts of DRP1 or OPA1, proteins involved in mitochondrial fission or fusion respectively (Figure 2D), a clear decrease in the mitochondrial localization of DRP1 was observed in irradiated cells, indicating that hyperfusion of the mitochondrial network was a consequence of a reduction in the levels of fission (Figure 2E-H). No difference was observed in the mitochondrial localization of OPA1 (Figure S3E). The specificity of the signal obtained with antibodies against DRP1 and OPA1 was evaluated by Western blot and immunofluorescence in cells transfected with siRNAs against DRP1 or OPA1 (Figure S3A-D).

To evaluate if the hyperfused mitochondrial network was electrically continuous, we performed microirradiation using a focalized laser in order induce depolarization of a small part of the network and evaluated the spread of depolarization through the mitochondrial network (Mitra et al., 2009). To do so, mitochondria were stained with both MitoTracker Green and TMRE and loss of TMRE was monitored both in the microirradiated region and in the entire mitochondrial network. We observed a fast depolarization of mitochondria within the microirradiated region as well as in areas of the network located at a distance of several micrometers from it, indicating connected mitochondrial branches (Figure 2I and S4). Interestingly, the mitochondrial membrane potential was rapidly recovered in the affected regions indicating the capacity of the mitochondrial network to compensate for localized damage.

Altogether, our results show a clear alteration of mitochondrial network induced by ionising radiation, with an increase in mtDNA levels, mitochondrial mass and membrane potential and hyperfusion of the mitochondrial network. All this adaptation responses, known to enhance cellular fitness and reduce apoptosis, could contribute to cellular radioresistance. Therefore, blocking this adaptive response by using small molecules targeting mitochondrial stability could have a major therapeutic potential for cancer treatment.

### G4 ligand RHPS4 localizes into mitochondria and blocks the irradiation induced mitochondrial adaptation

To assess if G4 ligands that stabilize mitochondrial G-quadruplexes can counteract mitochondrial adaptation to ionising radiation, we first treated cells with RHPS4, a G4 ligand that enters mitochondria and disrupts mtDNA transcription and replication, causing mitochondrial dysfunction (Falabella et al., 2018; Doimo et al., 2023). RHPS4 rapidly accumulated in the mitochondria of U2OS cells, as observed by live time imaging in cells in which the mitochondrial network was stained with MitoTracker Deep Red (Figure 3A, Figure S5). In agreement with previous observations (Falabella et al., 2018), no significant nuclear localization of RHPS4 was observed in living cells, suggesting that previously reported nuclear localization of RHPS4 is mostly due to the procedure of cell fixation resulting in a loss of mitochondrial membrane potential (Figure S5).

**Figure 3.**
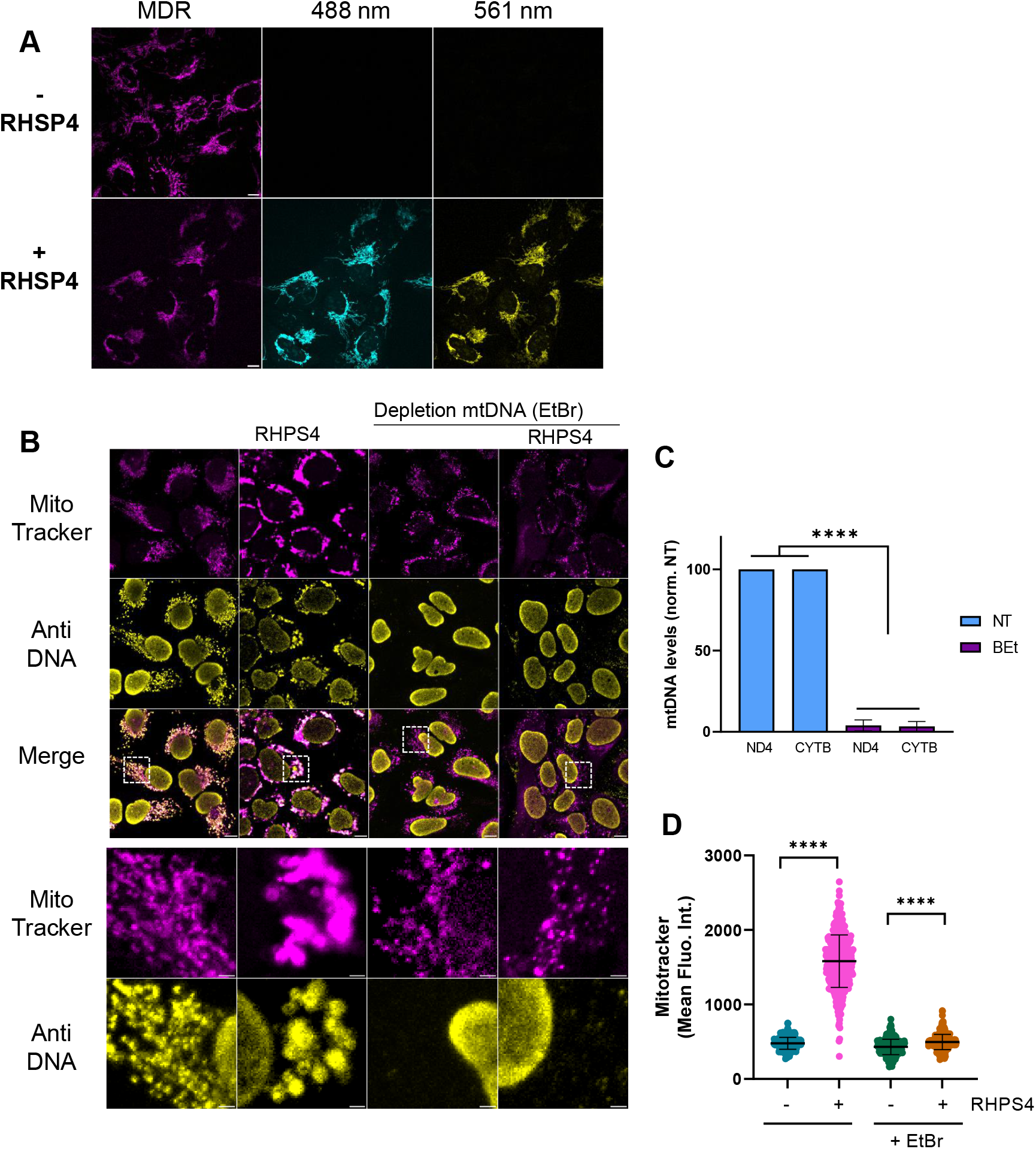
G4 ligand RHPS4 localizes to mitochondria and compromises the stability of mtDNA and mitochondrial morphology. **A)** U2OS cells were stained with MitoTracker Deep Red (magenta) and exposed to 10 µM of RHPS4 for 15 min before imaging with a spinning disk confocal microscope. RHPS4 fluorescence was imaged with 488nm (cyan) and 562nm (yellow) channels. **B)** U2OS cell treated or not with EtBr to deplete mtDNA were exposed to RHPS4 (10µM) for 24 H before staining with MitoTracker Red (magenta) and fixation. mtDNA was visualized with an anti-DNA antibody (yellow). Scale bar 10µm and 2µm for the zoom. **C)** Levels of mitochondrial DNA in control cells and in cells exposed to EtBr were measured by qPCR using probes against the mitochondrial genes ND4 and CYTB. Statistical analysis was performed with one-way ANOVA (****<0.0001). **D)** Mean intensity of MitoTracker Red was analyzed for more than 1500 cells for each condition from 3 independent experiments. Statistical analysis was performed in GraphPad using Kruskal-Wallis test (***P ≤ 0.001).

Exposure of cells to RHPS4 for 24 hours results in an alteration of mitochondrial morphology, with clustering of the mitochondria and nucleoids around the nucleus (Figure 3B-D). In order to evaluate if the effects of RHPS4 on mitochondrial morphology were dependent on the presence of mtDNA, we generated cells depleted of mtDNA by adding EtBr to the culture medium, inhibiting the replication of mtDNA. Loss of mtDNA was confirmed by qPCR (Figure 3D) and by immunofluorescence using an antibody against DNA (Figure 3B). No effect of RHPS4 on mitochondrial clustering was observed in cells depleted for mtDNA (Figure 3B,C), clearly indicating that RHPS4 induces mitochondrial dysfunction by directly targeting mtDNA.

The targeting of RHPS4 to mitochondria and its clear impact on mtDNA replication/loss and mitochondrial morphology, prompted us to evaluate if the addition of RHPS4 to the culture medium could limit the mitochondrial adaptive response induced by ionising radiation. For that, the levels of mtDNA were monitored at 48 and 96 hours after irradiation in the presence or absence of 0.5 µM of RHPS4. The ligand was added to the medium immediately before irradiation and kept all along the recovery period. The increase in mtDNA copy number induced by irradiation was clearly hampered in the presence of RHPS4, as measured both by qPCR (Figure 4A) and immunofluorescence analyses (Figure 4B-C). The same response was observed after irradiation with X-rays and in other tumoral cell lines such as T47D and HMLE (Figure S6A-C). Cytometry analysis showed that the addition of RHPS4 does not block the increase in cell size induced by irradiation (Figure 6D). However, the increase in mtDNA specifically observed In the cells that became bigger after irradiation is completely blocked in the presence of RHPS4 (Figure S6E), being not significantly different from the levels measured in non irradiated cells.

**Figure 4.**
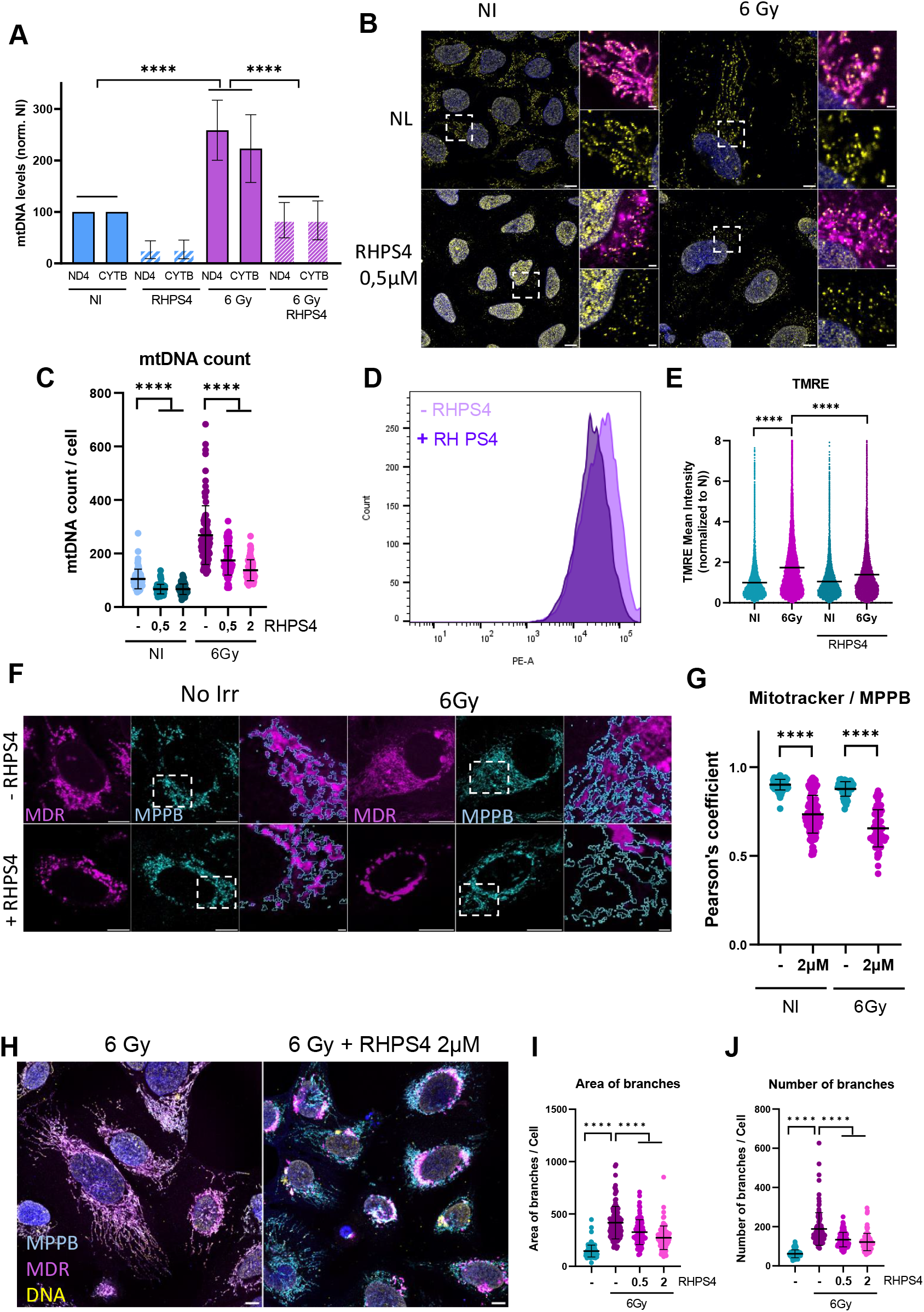
G4 ligand RHPS4 blocks the increases in mtDNA, mitochondrial mass and activity induced by ionising radiation. **A)** mtDNA copy number quantification by qPCR for two mtDNA regions, CYTB and ND4 48 or 96H after irradiation in the presence or absence of 0.5 µM RHPS4. Statistical analysis was performed for eight independent experiments with one-way ANOVA (****<0.0001). **B)** Confocal microscopy images of U2OS cells non-irradiated or 96H after irradiation, treated or not with RHPS4 (0,5µM). Cells were stained with MitoTracker Deep Red (magenta) before fixation and then stained with anti-DNA antibody (yellow) and DAPI (blue). Scale bar is 10µm and 2µm for zoom. **C)** Number of nucleoids was analysed for at least 100 cells for each condition from 4 independent experiments. **D)** TMRE probe was used to measure mitochondrial membrane potential by flow cytometry 4 days after irradiation (6Gy) in the presence or absence of RHPS4 (2µM). Histogram illustrates one representative experiments out of three. **E)** TMRE mean intensity from non-irradiated or irradiated cells with or without RHPS4 (2µM). 10.000 single cells were measured from 3 independent experiments and values normalised to NI cells. Statistical analysis were performed with GraphPad using Kruskal-Wallis test (***P ≤ 0.001). **F)** U2OS cells non irradiated or 96H after irradiation were stained with MitoTracker Deep Red (magenta) and anti-MPPB (cyan) and imaged by confocal microscopy. Scale bar 10 µm and 2 µm for the insets. **G)** Pearson correlation coefficient between MitoTracker Deep Red and MPPB in irradiated and non-irradiated cells in the absence or presence of RHPS4 (2µM). More than 50 cells from two independent experiments were analysed. **H)** Confocal microscopy images of irradiated U2OS cells 96H after irradiation, treated or not with 2 µM RHPS4. Cells were stained with MitoTracker Deep Red (cyan) before fixation and further stained with an anti-DNA (yellow) and anti-MPPB (magenta) antibodies; Scale bar is 10µm. **I)** Area and **J)** Number of mitochondrial branches per cell were analysed from 102 NI cells and 118 irradiated cells from 4 independent experiments. Statistical analysis was performed in GraphPad using Mann-Whitney test (***P ≤ 0.001).

In order to evaluate if irradiation, RHPS4 or their combination induced the accumulation of deletions or mutations at the level of the mtDNA molecule, sequence analysis of mtDNA was performed for all the different conditions 96H after irradiation. The use of the eKLIPse software, specifically developed to map mtDNA deletions (Goudenège et al., 2019) did not allow to detect accumulation of any particular deletion (Figure S7), in agreement with previous observations showing that RHPS4 efficiently blocks the progression of the mtDNA replication fork inducing the loss of the molecule (Doimo et al., 2023).

The increase of mitochondrial membrane potential measured by the TMRE probe observed in irradiated cells was also significantly reduced in cells irradiated in the presence of RHPS4 (Figure 4D-E). A lower mitochondrial membrane potential was also observed by microscopy in cells stained simultaneously with MitoTracker Deep Red, a dye accumulating in active mitochondria in a potential-dependent manner, and an antibody against MPPB allowing visualization of the mitochondrial matrix. While a perfect correlation was observed between both mitochondrial markers in the absence of RHPS4, a loss of MitoTracker Deep Red staining was observed in cells exposed to RHPS4, indicating a loss of mitochondrial membrane potential induced by the presence of RHPS4 in both irradiated and non-irradiated cells (Figure 4F-G).

In the same manner, hyperfusion of the mitochondrial network induced by irradiation was also affected in the presence of the ligand, with a clear reduction in the number of branches observed in cells exposed to irradiation in the presence of RHPS4 (Figure 4H-J).

### RHPS4 increases radiosensitivity in a mtDNA dependent manner

In agreement with previous reports (Berardinelli et al., 2015 and 2019), we observed a reduction in cellular proliferation and clonogenicity of U202 cells irradiated in the presence of RHPS4 (Figure 5A-C). The effect of the RHPS4 ligand in telomere maintenance has been proposed to be responsible for the radio-sensitizing effect of the molecule. However, we observed a reduction in cellular proliferation and survival after irradiation in the presence of very low concentrations of the ligand, from 0,5 to 2 µM, that have previously been shown not to induce detectable amount of nuclear DNA damage, as measured by γ-H2AX levels, or telomeric shortening (Fallabella et al., 2018). Therefore, in order to evaluate if the direct targeting of mtDNA by RHPS4 could was responsible for the increased sensitivity to irradiation, we performed a proliferation assay comparing cells depleted from mtDNA to normal cells. The depletion of mtDNA was induced by addition of ddC to the culture medium (Young et al., 2021). In our experimental conditions, a strong depletion of mtDNA levels was achieved after 5 days of exposure to ddC as measured both by immunofluorescence and qPCR (Figure 5D-F). While a reduction in cell survival was observed in WT cells irradiated in the presence of increasing amounts of RHPS4, the effect of the ligand was significantly reduced in cells depleted in mtDNA (Figure 5G), indicating that targeting the stability of mtDNA contributes to cellular radiosensitivity.

**Figure 5.**
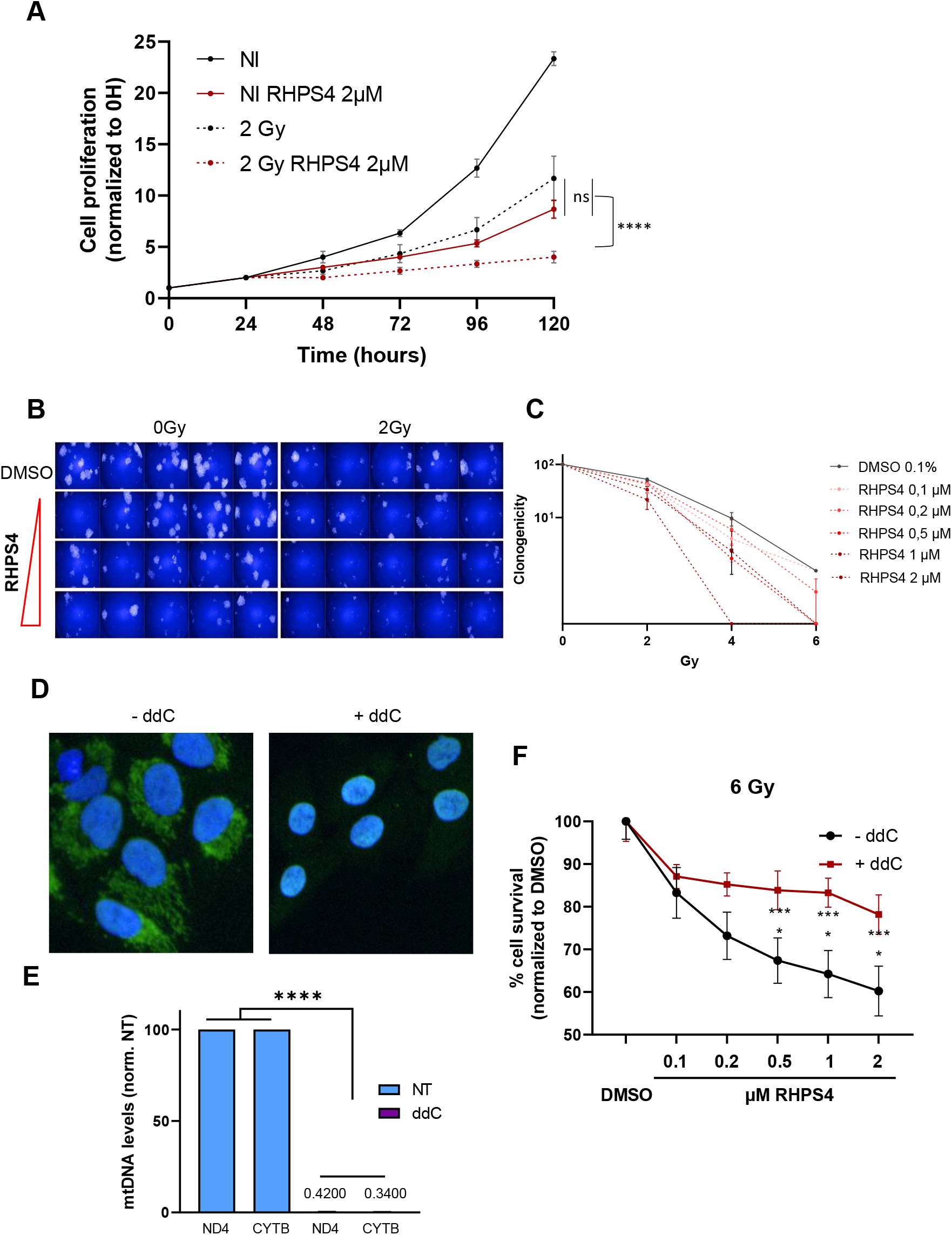
RHPS4 increases radiosensitivity in a mtDNA dependent manner. **A)** U2OS cell proliferation curves measured after a single irradiation at 2 Gy in the presence or absence of RHPS4 (2µM). Statistical analysis from three independent experiments was performed in GraphPad using Kruskal-Wallis test (n.s. not significant; **** p<0.0001). **B)** Images of a clonogenicity experiment performed at increasing concentrations of RHPS4 (0.5µM; 1µM and 2µM) combined or not with an irradiation at 2Gy. **C)** Clonogenicity measured at different irradiation doses (0; 2; 4 and 6 Gy) and different RHPS4 concentrations (0.1; 0.2; 0.5; 1; 2 µM). The number of clones were normalized to the number of clones obtained in the non irradiated cells fixed to 100%. Results are presented as a mean of three independent experiments. **D)** microscopy images of U2O2 cells depleted or not of their mtDNA by ddc treatment. mtDNA was stained with picogreen (green) and nuclear DNA with DAPI (blue). **E)** Quantification of mtDNA in cells treated or not by ddc was performed by qPCR using probes against CYTB and ND4. Statistical analysis was performed for five independent experiments with one-way ANOVA (****<0.0001). **G)** percentage of cell survival 6 days after an irradiation at 6Gy with increasing concentrations of RHPS4 doses (0.1; 0.2; 0.5; 1; 2 µM) in cells depleted (+ddC) or not (-ddC) for mtDNA. For E and G statistical analysis from three independent experiments was performed in GraphPad using Mann-Whitney test (***P ≤ 0.001).

**Figure 6.**
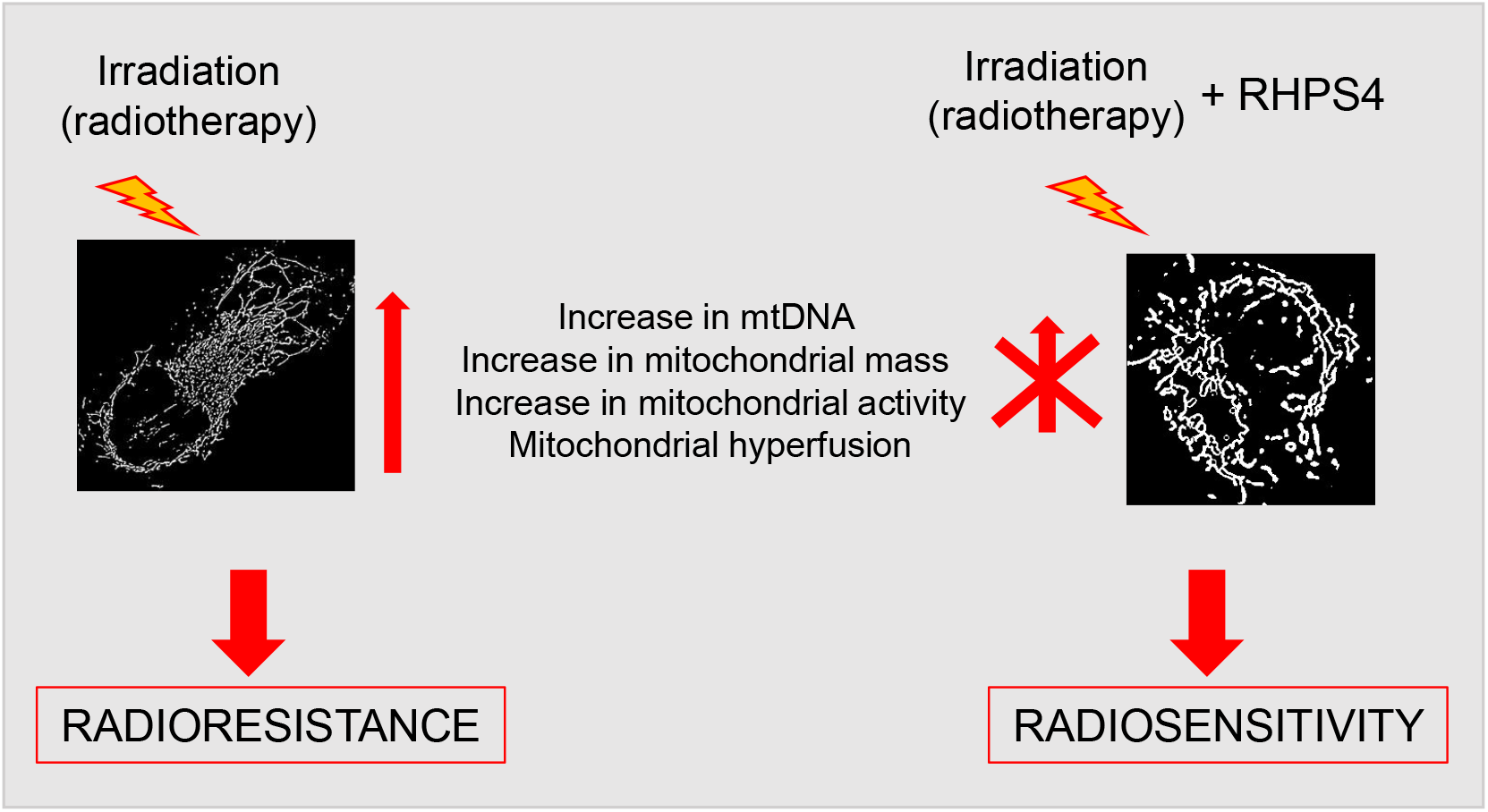
Radiosensitizing effects of RHPS4 mediated by mitochondrial targeting. RHPS4 interferes with the mitochondrial adaptation induced by ionising radiation by blocking the increases in mitochondrial DNA, mitochondrial mass, activity and the hyperfusion of the network, therefore reducing cellular radioresistance.

## DISCUSSION

Our results show that cells surviving ionizing radiation develop a mitochondrial adaptation resulting in an increase in mtDNA, mitochondrial mass and membrane potential (Figure 1). An increase in mtDNA copy number and mitochondrial biogenesis is considered to be a compensatory mechanism aimed at sustaining OXHPOS activity in order to overcome the eventual bioenergetics defects induced by mtDNA damage to some molecules (Filograna et al., 2019). Indeed, several studies have demonstrated that increasing mtDNA copy number by different genetic tools, boosts mitochondrial biogenesis and rescues mitochondrial dysfunction and may provide a novel therapeutic strategy for the treatment of mitochondrial diseases (Filograna et al., 2021).

The irradiation-induced increase in mitochondrial mass is correlated with an mitochondrial hyperfusion. Mitochondrial hyperfusion is a cellular response to various stress conditions, characterized by the extensive merging of mitochondria into elongated and interconnected networks. The so-called Stress induced mitochondrial hyperfusion (SIMH) was described for the first time in 2009 as an adaptative pro-survival response of the cells against stress (Tondera et al., 2009). Mitochondrial hyperfusion has been observed after UV exposure, cycloheximide and Actinomycin D treatments, or serum deprivation. Although to our knowledge no previous reports have shown ionising radiation induced SIMH, a slight increase in mitochondrial elongation, together with an increase in respiration, following IR has been recently reported in irradiated mesenchymal stem cells (Patten et al., 2019). Mitochondrial hyperfusion results from a disequilibrium between fusion and fission and is greatly reduced in cells deficient in MFN1/2 and OPA1, required for the fusion of outer and inner mitochondrial membrane respectively (Westermann, 2012). The decrease mitochondrial DRP1, a key regulator of mitochondrial fission, we observed in irradiated cells (Figure 2) suggests that a reduced fission is at the origin of the observed hyperfusion of the network. These results are in agreement with previous publications showing that phosphorylation of Drp1-S637 is required for mitochondrial elongation after starvation (Gomes et al., 2011). The results presented in Figure 3 and Figure S4 show that the hyperfused mitochondrial networks is electrically continuous. Therefore, mitochondrial damage induced in a small part of the network by laser microirradiation, rapidly spreads affecting other regions that are a several µm away. This electrical continuity has been shown to enhance the overall bioenergetic efficiency of the mitochondria, allowing the cell to optimize ATP production under stress conditions (Westermann., 2012; Mitra et al., 2009; De Giorgi et al., 2000). The interconnected network can therefore protect against localized damage, as healthy mitochondria can support and compensate for the impaired ones within the network, as can be observed by the fast recovery of TMRE in regions affected by microirradiation induced damage (Figure 3 and S4).

The adaptive mitochondrial response could help cells to better cope with and recover from stressful conditions, supporting overall cellular survival and radioresistance. Recent findings highlight the central role of mitochondrial adaptation in chemoresistance of acute myeloid leukemia (AML). An increase in mitochondrial mass and respiratory activity has been observed in cells developing drug resistance and several molecules targeting mitochondrial metabolism or dynamics have been shown to enhance cellular sensitivity to different chemotherapeutical agents and open new therapeutical avenues for AML (Bosc et al., 2021; Larrue et al., 2023).

Mitochondria are essential for the proliferation of cancer cells, not only because of their main role in ATP production but also because of their essential roles in cellular signaling, apoptosis and pyrimidine synthesis needed for DNA replication. Therefore, many molecules inducing a mitochondrial dysfunction are gaining interest for their potential use in cancer treatment. The term MITOCANS, for MITOchondria and CANcer, has even been proposed for anti-cancer drugs that induce mitochondrial dysfunction by targeting different mitochondrial pathways (Neuzil et al., 2020). mtDNA being essential for OXPHOS biogenesis, several molecules targeting mtDNA stability, both indirectly by inducing mtDNA damage and dysfunction or directly by binding or cleaving mtDNA are emerging as promising tools for cancer treatment (Lin et al., 2023; Lin et al., 2023; Mukherjee et al., 2023). Among the proteins regulating mtDNA transcription and replication, TFAM and POLRMT play an essential role and have been identified as potential targets for anti-cancer therapy. The anti-tumor efficacy of POLRMT inhibitors has been confirmed in ovarian, lung and cervical cancer (Bonekamp et al., 2020). Considering the recent discovery of a direct link between high mtDNA levels and accelerated lung cancer progression, along with the fact that impaired mtDNA copy number increase hinders tumor growth, targeting mtDNA offers an interesting perspective for cancer treatment (Mennuni et al., 2024).

In this context, targeting mitochondrial G4 by using mitochondrial localized G4 ligands with the capacity to block transcription and replication of mtDNA (Fallabella et al., 2019) represents a promising therapeutical strategy. Up to now, radiosensitizing effects of G4 ligands, including RHPS4, have been explained by their effect on telomere stability, replicative stress and cell cycle arrest (Berardinelli et al., 2015 and 2019). Interestingly, a systematic comparison of the effects of the RHPS4 at different concentrations has shown that while telomeric dysfunction and accumulation of nuclear DSB are observed at higher concentrations of the ligand (10 µM), the incubation of cells with a low dose (2 µM) interferes with mtDNA copy number but does not induce direct detectable nuclear DNA damage (Falabella et al., 2019). In agreement with previous findings, we have observed that RHPS4 preferentially localizes to mitochondria in living cells and affects mtDNA content and mitochondrial morphology by directly targeting mtDNA (Figure 3A, S5; Falabella et al., 2019). Therefore, an alternative, though not exclusive, explanation for the telomere instability induced by RHPS4 could involve an indirect effect resulting from mitochondrial dysfunction. Interestingly, oxidative stress has been shown to accelerate telomere loss (von Zglinicki, 2002) and a direct and tight link between mitochondrial dysfunction and telomere instability has been stablished. The use of a chemoptogenetic tool allowing the induction of singlet oxygen specifically in the mitochondrial matrix, has recently been show to induce a secondary wave of superoxide and hydrogen peroxide and rapid telomere dysfunction as characterized by 53BP1-positive TIFs, telomere fragility, and telomere loss in the apparent absence of general nuclear DNA strand breaks (Qian et al., 2019). These findings could explain the telomere instability observed in cells exposed to low concentrations of RHPS4 (0.2, 0.5 or 1µM) during 5 days (Berardinelli et al., 2019), that could indeed be indirectly induced by a mitochondrial dysfunction. Altogether those results suggest that mtDNA could be the primary target of RHPS4. Similar results have been obtained with G4 ligands derived from BMVC that accumulate in mitochondria of cancer cells and can induce cancer cell death without damaging normal cells. Interestingly, an in agreement with our observations showing that radiosensitizig effects of RHPS4 are dependent on mtDNA, no effect of the ligand on cell death was observed in cells depleted for mtDNA, indicating that mtDNA is the major target of the G4 ligand BMVC (Huang et al., 2015).

Our results show that the G4 ligand RHPS4, with a primary mitochondrial localization, specifically target mtDNA and induce mtDNA loss and mitochondrial dysfunction resulting in a blockage of the adaptative mitochondrial response induced by ionising irradiation and therefore increasing radiosensitivity. These results are not only a major fundamental advance concerning the understanding of radiation effects on mitochondrial network and the formation of G4 in mtDNA but also offer new therapeutical possibilities for the use of this molecule and other mitochondrial imported G4 ligands in cancer treatment.

## MATERIAL AND METHODS

### Cell culture

All cell lines were cultivated in Heraeus Thermo Scientific BBD6220 incubator at 37°C in a humidified atmosphere of 5% CO2. Absence of mycoplasma was regularly tested with the MycoAlert™ Mycoplasma detection kit from Lonza Group, Ltd. U2OS cells were purchased from ATCC, tumor breast cancer cell lines T47D were obtained from the American Type Culture Collection (Rockville, MD) and Human Mammary Epithelial HMLE cell line was kindly provided by Professor Robert A. Weinberg (Whitehead Institute, Cambridge, MA, USA). All cell lines were grown at 37°C in adherent conditions in cell culture media supplemented with 10% fetal bovine serum (FBS) (Sigma, F7524) and 5% penicillin-streptomycin (Gibco 15140122). U2OS were cultured in DMEM, low glucose, GlutaMAX™ Supplement, pyruvate (Gibco; Thermo Fisher Scientific, Inc.). mtDNA depletion was induced by incubating cells in DMEM High glucose containing 100 ng/ml BEt or 10µM of 2,3 dideoxycitidine (ddC) complemented with 50 µg/ml Uridine. For T47D and MCF7, Dulbecco’s Modified Eagle’s Medium (DMEM), high glucose, GlutaMAX™ Supplement, pyruvate (Gibco; Thermo Fisher Scientific, Inc.) was used. HMLE cells were cultured in DMEM/F-12 Nutrient Mixture, GlutaMAX Supplement (Gibco; Thermo Fisher Scientific, Inc.) with 10 ng/ml human epidermal growth factor, 0.5 μg/ml hydrocortisone and 10 μg/ml insulin (all from Sigma-Aldrich, Merck).

### Cell treatments and irradiations

Gamma-irradiations were performed on a GSR D1 irradiator (Gamma Medical Service, Leipzig, Germany) and cells harvested a. This self-shielded device irradiates with four sources of 137Cs, with a total activity around 180.28 TBq (measured in March 2014). The samples were irradiated at different single doses of 2, 4, or 6 Gy, with a dose rate of 1.5 Gy/min, taking the radioactive decrease into account. The samples were irradiated in 6 well plates or Ibidi µ-Slide 4-well plates and recovered at the indicated times after irradiation. Prior to irradiation, dosimetry was performed. A cylindrical ionizing chamber 31010 by PTW was used as the recommendation of the AAPM’S TG-61. This ionizing chamber has a cavity of 0.125 cm3 calibrated in 137Cs air kerma free in air at the PTB reference facility number 1904442 and 2108382401. The polarity and the ion recombination were measured for this 137Cs source. Each measurement was corrected by the KTP factor to take the variation of temperature and atmospheric pressure into account.

For X-rays irradiation, an irradiator ELEKTA VERSA HD with a 6 MV photon beam was used with a dose rate of 4.5 Gy/min and an irradiation field size of 30 × 30 cm^2^. The irradiated plates were positioned at the treatment isocenter, which corresponds to a source-to-surface distance (SSD) of 1 meter. At this point, the dose and dose rate were determined. Before reaching the plates, the beam passed through 8 cm of PMMA (polymethyl methacrylate), followed by an additional 2 cm. The delivered dose was measured prior to each irradiation series using a Farmer ionization chamber (PTW 30013), calibrated in water-equivalent conditions. The chamber was placed in the same position and under the same conditions as the plates to ensure accurate dose and dose rate measurements.

RHPS4 ligand (RHPS 4-methosulfate; Tocris; Ref 5311) was added to the culture medium at the indicated concentrations for 30 min before irradiation and kept in the medium for all the recovery period before cell harvest or fixation.

### qPCR

Genomic DNA was extracted from 1 million cells using the Qiamp DNA mini kit (QIAGEN, #51304) following manufacturer instructions and DNA concentration was measured with Nanodrop 2000c (Thermo Fisher Scientific). 75ng of DNA was used for qPCR with the iTaq Universal Probes Supermix (Biorad #1725130) using the Applied Biosystems™ QuantStudio™ three Real-Time PCR System. Taqman probes against ND1 (Hs02596873_s1), ND4 (Hs02596876_g1), COX2 (Hs02596865_g1), COX3 (Hs02596866_g1), CYTB (Hs02596867_s1), tRNA1 (Hs02596871_s1) and tRNA2 (Hs02596871_s1) were purchased from Thermo Fisher Scientific. Each sample was run in triplicate. Mean Ct values were used to calculate the mtDNA copy number relative to untreated samples using the ΔΔCt method.

### Western blot

Cell pellets (from 3 × 10^6^ cells) were resuspended in 50 µl of Lysis buffer (Tris 20 mM, NaCl 20 mM, SDS 0.1%, MgCl2 1 Mm, Benzonase 0.25 U/µl and Protease inhibitors cocktail 1X) and sonicated for 10 min (with pulses 30s on/30s off). After sonication the samples were centrifuged at 12000 at 4°C for 5 min and the supernatants were recovered. Protein concentration was measured with the Bradford assay (Bio-Rad #500-0006), Laemmli buffer was added at 1x concentration (0.1% 2-mercaptoethanol, 0.0005% Bromophenol blue, 10% glycerol, 2% SDS, and 63 mM Tris pH 6.8) before heating for 5 min at 95°C, and 20 µg of protein extracts were loaded in a mini PROTEAN TGX stain-free gel (Bio-Rad #4568083). Precision plus protein dual color standards (Bio-Rad #1610374) was used as a Molecular weight marker. The transfer on nitrocellulose membrane (Bio-Rad #1704157) was performed with the Trans-Blot® Turbo™ Transfer System (Bio-Rad #1704150). The membrane was blocked for 1 h in blocking solution (PBS-0.1% Tween20 containing 5% milk), and incubated for 1 h at room temperature with primary antibodies (Table S1). Membranes were washed three times for 5 min with PBS-0.1% Tween20 and further incubated for 45 min in secondary antibodies anti-rabbit IR800 (Diagomics R-05060) and anti-mouse IR700 (Diagomics R-05055) diluted at 1/10.000 in PBS-0.1% Tween 20 containing 5% milk. Western blots were imaged with the Li-Cor Odyssey DLx system

### Flow cytometry, acquisition and analysis

Mitochondrial mass and mitochondrial membrane potential were analyzed using MitoTracker Green (Invitrogen; Thermo Fisher Scientific M7513) and TMRE (Sigma-Aldrich 87917) respectively. Following soft trypsinization, cells were loaded with 200 nM MitoTracker Green and/or 10 nM TMRE and incubated for 20 min at 37°C, then immediately analyzed by flow cytometry. Cells were analyzed on a SORP LSR-II analyzer (configuration: 488, 561, 405, 355 and 635 nm; BD Biosciences) or BD FACSCalibur (configuration: 488 and 635 nm; BD Biosciences). Data were analyzed with FlowJo v10.7.1 (Tree Star).

### Immunofluorescence, live cell microscopy and image analysis

For microscopy experiments cells were grown on Ibidi µ-Slide 4-well plates for two days before experiment. For live imaging cells were washed twice with warm DMEM and incubated with 100nM of MitoTracker (Green, Red or Deep Red as indicated), or 10 nM TMRE (Invitrogen; Thermo Fisher Scientific, Inc.) for 20min at 37°C followed by washing with complete medium. Image acquisition was performed using a NIKON Ti2 / GATACA, W1 spinning disk microscope with 60X oil immersion objective (Plan Apo N.A. 1.4) in a chamber maintained at 37°C with 95% relative humidity, 5% CO2 and 18% oxygen).

For Immunofluorescence experiments, cells were washed with DMEM and immediately fixed for 20min at 37°C with 2% formaldehyde in DMEM solution, rinsed with PBS and permeabilised at room temperature in PBS 0.1% Triton (X100) for 5 min. Cells were incubated in blocking solution (PBS containing 0.1% Triton 100X, 3% BSA and 1% normal goat serum) at 37°C for 1h. Cells were incubated with primary antibodies (Table S1) diluted in blocking solution for 1h at 37°C, washed three times for 5 min in PBS containing 0.1% Triton and further incubated with secondary antibodies (Table S1) diluted 1/1000 in blocking solution for 45 min at 37°C. Cells were washed three times for 5 min in PBS containing 0.1% Triton. Nuclear DNA was counterstained with 1 µg/ml 4′,6′-diamidino-2-phenylindole (DAPI).

Image acquisition was performed using a NIKON Ti2 / GATACA, W1 spinning disk microscope with 60X oil immersion objective (Plan Apo N.A. 1.4) or a Leica SP8 confocal microscope with a 63× oil immersion objective (HC PL Apo N.A. 1.4). Image analysis was performed with ImageJ/Fiji (Schneider et al., 2012). Image analysis was performed with ImageJ. Mitochondrial morphology alterations were analysed by applying the Kernel algorithm (Koopman et al., 2005) to binary images. This approach allows measurement of morphological parameters for individual mitochondria, such as a measure of mitochondrial count and area. Quantification of the number of mitochondrial nucleoids was performed with an anti-DNA after subtraction of the nucleus stained with DAPI. Pearson and Manders correlation coefficients between were calculated with ImageJ using the plugin JACoP (Bolte and Cordelieres, 2006).

For each parameter, the number of cells analysed from at least two or three independent experiments is indicated in the legend to the figures.

### Sequencing and analysis of mtDNA

The entire mtDNA was amplified in overlapping fragments using a combination of primers designed to avoid nuclear pseudogene amplification and already previously validated in Rho zero cells devoid of mtDNA. Primer sequences, polymerase chain reaction (PCR) conditions, and sequencing were performed as previously described, and data analyzed with the dedicated software eKLIPse, a sensitive and specific tool based on soft clipping, allowing the univocal detection of mtDNA rearrangements and breakpoints, and their quantification from NGS data (Goudenège et al., 2019).

### Proliferation/survival assay

When required, U2OS cells were pretreated with 10µM of 2,3 dideoxycitidine (ddC) for 3 days before the plate seeding, and then throughout the experiment, to induce and maintain mtDNA depletion. Cells – either mtDNA depleted or not - were seeded in black-walled, clear-bottom 96-well culture plates (Costar #3904, Corning). For endpoint measurement 6 days post-seeding in non-irradiated and irradiated conditions, 300 cells were plated per well. For daily measurement of cell growth in non-irradiated and irradiated conditions, 750 and 2000 cells respectively were plated per well. Cells were treated with RHPS4 the same day (for 6-days experiments) or one day post-seeding (for daily growth measurements) at concentrations ranging from 100 nM to 2 µM, or with DMSO as a vehicle control and irradiated 3 hours later. Plates were incubated at 37°C, then fixed at the indicated timepoints post-RHPS4 treatments with 2% (w/v) paraformaldehyde. Nuclear DNA was stained with 2 µg/mL Hoechst 33342 (Sigma). Images were captured using an automated epifluorescence microscope (Operetta, Perkin Elmer) at 10× magnification, with 9 fields per well in the blue fluorescence channel (excitation: 380 ± 20 nm, emission: 445 ± 35 nm). An automated algorithm, developed in Harmony 3.0 (Perkin Elmer), was used to quantify cell numbers. In brief, nuclear regions of interest (ROIs) were segmented based on Hoechst staining, and the relative cell proliferation/survival in each condition was calculated as the number of cells in experimental conditions relative to the average cell count in the reference control wells.

### Clonogenicity assay

U2OS cells were seeded in black-walled, clear-bottom 96-well culture plates (Costar #3904, Corning) at densities of 15 cells/well. Cells were then treated with RHPS4 the same day at concentrations ranging from 100 nM to 2 µM, or with DMSO as a vehicle control and irradiated 3 hours later. Plates were incubated at 37°C, then fixed at the indicated timepoints post-RHPS4 treatments with 2% (w/v) paraformaldehyde, and nuclear DNA was stained with 4 µg/mL Hoechst 33342 (Sigma). Images of clones were captured using an automated epifluorescence microscope (Operetta, Perkin Elmer) at 2× magnification, with 1 field per well in the blue fluorescence channel (excitation: 380 ± 20 nm, emission: 445 ± 35 nm). An automated algorithm, developed in Harmony 3.0 (Perkin Elmer), was used to quantify clone numbers. In brief, nuclear regions of interest (ROIs) were segmented as fluorescent spots based on Hoechst staining. Clones were then defined as clusters of nuclei in close proximity (4 pixels or less) and made of 50 cells or more. For each condition, clone amount represents the sum of clones in ten wells. The relative clone amount was calculated as the number of clones in the experimental conditions relative to the average clone amount in the reference control wells.

### Statistical analysis

Statistical analysis was performed using GraphPad. The statistical test used and the number of cells and independent experiments analyzed are indicated in the legend to the figures.

## AUTHOR CONTRIBUTION

S.T. performed experiments and analysis and prepared the figures; C.D., I.B., C.S. J.L., performed experiments and analysis. G.P., G.K., M.V., performed proliferation and clonogenicity experiments and analysis. G.L. performed sequence analysis. J.P.R. obtained funding, read and edited the manuscript. A. C. conceptualized and supervised the project, performed experiments and analysis, prepared figures, wrote the manuscript and obtained funding.

## ACKNOWLEDGEMENTS

We would like to thank Arturo Londono, Alain Nicolas and Marie-Paule Teulade-Fichou for fruitful discussions all along the project, Xavier Renaudin for critical reading of the manuscript, Véronique Ménard, Mathieu Agelou and Guillaume Boissonnat for their help in irradiation experiments. We thank the Ligue contre le cancer, CEA radiobiology program, and Electricité de France for the financial support.

## REFERENCES

Begg, A.C., Stewart, F.A. and Vens, C. (2011) Strategies to improve radiotherapy with targeted drugs. Nat. Rev. Cancer, 11, 239–253.

Berardinelli, F., Siteni, S., Tanzarella, C., Stevens, M.F., Sgura, A. and Antoccia, A. (2015) The G-quadruplex-stabilising agent RHPS4 induces telomeric dysfunction and enhances radiosensitivity in glioblastoma cells. DNA Repair, 25, 104–115.

Berardinelli, F., Tanori, M., Muoio, D., Buccarelli, M., Di Masi, A., Leone, S., Ricci-Vitiani, L., Pallini, R., Mancuso, M. and Antoccia, A. (2019) G-quadruplex ligand RHPS4 radiosensitizes glioblastoma xenograft in vivo through a differential targeting of bulky differentiated- and stem-cancer cells. J. Exp. Clin. Cancer Res., 38, 311.

Bharti, S.K., Sommers, J.A., Zhou, J., Kaplan, D.L., Spelbrink, J.N., Mergny, J.-L. and Brosh, R.M. (2014) DNA Sequences Proximal to Human Mitochondrial DNA Deletion Breakpoints Prevalent in Human Disease Form G-quadruplexes, a Class of DNA Structures Inefficiently Unwound by the Mitochondrial Replicative Twinkle Helicase. J. Biol. Chem., 289, 29975–29993.

Bonekamp, N.A., Peter, B., Hillen, H.S., Felser, A., Bergbrede, T., Choidas, A., Horn, M., Unger, A., Di Lucrezia, R., Atanassov, I., et al. (2020) Small-molecule inhibitors of human mitochondrial DNA transcription. Nature, 588, 712–716.

Bosc, C., Saland, E., Bousard, A., Gadaud, N., Sabatier, M., Cognet, G., Farge, T., Boet, E., Gotanègre, M., Aroua, N., et al. (2021) Mitochondrial inhibitors circumvent adaptive resistance to venetoclax and cytarabine combination therapy in acute myeloid leukemia. Nat. Cancer, 2, 1204–1223.

Chen, Y., Gao, H. and Ye, W. (2018) Mitochondrial DNA Mutations Induced by Carbon Ions Radiation: A Preliminary Study. Dose-Response, 16, 1559325818789842.

Dahal, S., Siddiqua, H., Katapadi, V.K., Iyer, D. and Raghavan, S.C. (2022) Characterization of G4 DNA formation in mitochondrial DNA and their potential role in mitochondrial genome instability. FEBS J., 289, 163–182.

De Giorgi, F., Lartigue, L. and Ichas, F. (2000) Electrical coupling and plasticity of the mitochondrial network. Cell Calcium, 28, 365–370.

Doimo, M., Chaudhari, N., Abrahamsson, S., L’Hôte, V., Nguyen, T.V.H., Berner, A., Ndi, M., Abrahamsson, A., Das, R.N., Aasumets, K., et al. (2023) Enhanced mitochondrial G-quadruplex formation impedes replication fork progression leading to mtDNA loss in human cells. Nucleic Acids Res., 51, 7392–7408.

Dong, L., Gopalan, V., Holland, O. and Neuzil, J. (2020) Mitocans Revisited: Mitochondrial Targeting as Efficient Anti-Cancer Therapy. Int. J. Mol. Sci., 21, 7941.

Falabella, M., Kolesar, J.E., Wallace, C., De Jesus, D., Sun, L., Taguchi, Y.V., Wang, C., Wang, T., Xiang, I.M., Alder, J.K., et al. (2019) G-quadruplex dynamics contribute to regulation of mitochondrial gene expression. Sci. Rep., 9, 5605.

Figueiredo, J., Mergny, J.-L. and Cruz, C. (2024) G-quadruplex ligands in cancer therapy: Progress, challenges, and clinical perspectives. Life Sci., 340, 122481.

Filograna, R., Koolmeister, C., Upadhyay, M., Pajak, A., Clemente, P., Wibom, R., Simard, M.L., Wredenberg, A., Freyer, C., Stewart, J.B., et al. (2019) Modulation of mtDNA copy number ameliorates the pathological consequences of a heteroplasmic mtDNA mutation in the mouse. Sci. Adv., 5, eaav9824.

Filograna, R., Mennuni, M., Alsina, D. and Larsson, N. (2021) Mitochondrial DNA copy number in human disease: the more the better? FEBS Lett., 595, 976–1002.

Gomes, L.C., Benedetto, G.D. and Scorrano, L. (2011) During autophagy mitochondria elongate, are spared from degradation and sustain cell viability. Nat. Cell Biol., 13, 589–598.

Goudenège, D., Bris, C., Hoffmann, V., Desquiret-Dumas, V., Jardel, C., Rucheton, B., Bannwarth, S., Paquis-Flucklinger, V., Lebre, A.S., Colin, E., et al. (2019) eKLIPse: a sensitive tool for the detection and quantification of mitochondrial DNA deletions from next-generation sequencing data. Genet. Med., 21, 1407–1416.

Huang, W.-C., Tseng, T.-Y., Chen, Y.-T., Chang, C.-C., Wang, Z.-F., Wang, C.-L., Hsu, T.-N., Li, P.-T., Chen, C.-T., Lin, J.-J., et al. (2015) Direct evidence of mitochondrial G-quadruplex DNA by using fluorescent anti-cancer agents. Nucleic Acids Res., 10.1093/nar/gkv1061.

Jin, X., Li, F., Liu, B., Zheng, X., Li, H., Ye, F., Chen, W., Li, A. (2018) Different mitochondrial fragmentation after irradiation with X-rays and carbon ions in HeLa cells and its influence on cellular apoptosis. Biochemical and Biophysical Research Communications, 500(4), 958–965.

Larrue, C., Mouche, S., Lin, S., Simonetta, F., Scheidegger, N.K., Poulain, L., Birsen, R., Sarry, J.-E., Stegmaier, K. and Tamburini, J. (2023) Mitochondrial fusion is a therapeutic vulnerability of acute myeloid leukemia. Leukemia, 37, 765–775.

Lin, Y., Yang, B., Huang, Y., Zhang, Y., Jiang, Y., Ma, L. and Shen, Y.-Q. (2023) Mitochondrial DNA-targeted therapy: A novel approach to combat cancer. Cell Insight, 2, 100113.

Liu, Y., Sun, Y., Guo, Y., Shi, X., Chen, X., Feng, W., Wu, L.-L., Zhang, J., Yu, S., Wang, Y., et al. (2023) An Overview: The Diversified Role of Mitochondria in Cancer Metabolism. Int. J. Biol. Sci., 19, 897–915.

McCann, E., O’Sullivan, J. and Marcone, S. (2021) Targeting cancer-cell mitochondria and metabolism to improve radiotherapy response. Transl. Oncol., 14, 100905.

Mennuni, M., Wilkie, S.E, Michon, P., Alsina, D., Filograna, R., Lindberg, M., Sanin, D.E. et al. (2024) High Mitochondrial DNA Levels Accelerate Lung Adenocarcinoma Progression. Science Advances 10 (44).

Merle, P., Gueugneau, M., Teulade-Fichou, M.-P., Müller-Barthélémy, M., Amiard, S., Chautard, E., Guetta, C., Dedieu, V., Communal, Y., Mergny, J.-L., et al. (2015) Highly efficient radiosensitization of human glioblastoma and lung cancer cells by a G-quadruplex DNA binding compound. Sci. Rep., 5, 16255.

Mitra, K., Wunder, C., Roysam, B., Lin, G. and Lippincott-Schwartz, J. (2009) A hyperfused mitochondrial state achieved at G 1 –S regulates cyclin E buildup and entry into S phase. Proc. Natl. Acad. Sci., 106, 11960–11965.

Mukherjee, S., Bhatti, G.K., Chhabra, R., Reddy, P.H. and Bhatti, J.S. (2023) Targeting mitochondria as a potential therapeutic strategy against chemoresistance in cancer. Biomed. Pharmacother., 160, 114398.

Neuzil, J., Dong, L.-F., Rohlena, J., Truksa, J. and Ralph, S.J. (2013) Classification of mitocans, anti-cancer drugs acting on mitochondria. Mitochondrion, 13, 199–208.

Prithivirajsingh, S., Story, M.D., Bergh, S.A., Geara, F.B., Kian Ang, K., Ismail, S.M., Stevens, C.W., Buchholz, T.A. and Brock, W.A. (2004) Accumulation of the common mitochondrial DNA deletion induced by ionising radiation. FEBS Lett., 571, 227–232.

Qian, W., Kumar, N., Roginskaya, V., Fouquerel, E., Opresko, P.L., Shiva, S., Watkins, S.C., Kolodieznyi, D., Bruchez, M.P. and Van Houten, B. (2019) Chemoptogenetic damage to mitochondria causes rapid telomere dysfunction. Proc. Natl. Acad. Sci., 116, 18435–18444.

Sahayasheela, V.J., Yu, Z., Hidaka, T., Pandian, G.N. and Sugiyama, H. (2023) Mitochondria and G-quadruplex evolution: an intertwined relationship. Trends Genet., 39, 15–30.

Schilling-Tóth, B., Sándor, N., Kis, E., Kadhim, M., Sáfrány, G. and Hegyesi, H. (2011) Analysis of the common deletions in the mitochondrial DNA is a sensitive biomarker detecting direct and non-targeted cellular effects of low dose ionising radiation. Mutat. Res. Mol. Mech. Mutagen., 716, 33–39.

Tan, A.S., Baty, J.W., Dong, L.-F., Bezawork-Geleta, A., Endaya, B., Goodwin, J., Bajzikova, M., Kovarova, J., Peterka, M., Yan, B., et al. (2015) Mitochondrial Genome Acquisition Restores Respiratory Function and Tumorigenic Potential of Cancer Cells without Mitochondrial DNA. Cell Metab., 21, 81–94.

Tilokani, L., Nagashima, S., Paupe, V. and Prudent, J. (2018) Mitochondrial dynamics: overview of molecular mechanisms. Essays Biochem., 62, 341–360.

Tondera, D., Grandemange, S., Jourdain, A., Karbowski, M., Mattenberger, Y., Herzig, S., Da Cruz, S., Clerc, P., Raschke, I., Merkwirth, C., et al. (2009) SLP-2 is required for stress-induced mitochondrial hyperfusion. EMBO J., 28, 1589–1600.

Von Zglinicki, T. (2002) Oxidative stress shortens telomeres. Trends Biochem. Sci., 27, 339–344.

Wallace, D.C. (2012) Mitochondria and cancer. Nat. Rev. Cancer, 12, 685–698.

Wanrooij, P.H., Uhler, J.P., Shi, Y., Westerlund, F., Falkenberg, M. and Gustafsson, C.M. (2012) A hybrid G-quadruplex structure formed between RNA and DNA explains the extraordinary stability of the mitochondrial R-loop. Nucleic Acids Res., 40, 10334–10344.

Wanrooij, P.H., Uhler, J.P., Simonsson, T., Falkenberg, M. and Gustafsson, C.M. (2010) G-quadruplex structures in RNA stimulate mitochondrial transcription termination and primer formation. Proc. Natl. Acad. Sci., 107, 16072–16077.

Westermann, B. (2012) Bioenergetic role of mitochondrial fusion and fission. Biochim. Biophys. Acta BBA - Bioenerg., 1817, 1833–1838.

Youle, R.J. and Van Der Bliek, A.M. (2012) Mitochondrial Fission, Fusion, and Stress. Science, 337, 1062–1065.

Young, C.K.J., Wheeler, J.H., Rahman, Md.M. and Young, M.J. (2021) The antiretroviral 2′,3′-dideoxycytidine causes mitochondrial dysfunction in proliferating and differentiated HepaRG human cell cultures. J. Biol. Chem., 296, 100206.

Yuan, Y., Ju, Y.S., Kim, Y., Li, J., Wang, Y., Yoon, C.J., Yang, Y., Martincorena, I., Creighton, C.J., Weinstein, J.N., et al. (2020) Comprehensive molecular characterization of mitochondrial genomes in human cancers. Nat. Genet., 52, 342–352.

Yusoff, A.A.M., Wan Abdullah, W.S., Mohd Khair, S.Z.N. and Abd Radzak, S.M. (2019) A comprehensive overview of mitochondrial DNA 4977-bp deletion in cancer studies. Oncol. Rev., 13.

Zaffaroni, M., Vincini, M.G., Corrao, G., Marvaso, G., Pepa, M., Viglietto, G., Amodio, N. and Jereczek-Fossa, B.A. (2022) Unraveling Mitochondrial Determinants of Tumor Response to Radiation Therapy. Int. J. Mol. Sci., 23, 11343.

