## Supplementary Figures for "G-quadruplex ligand RHPS4 compromises cellular radio-resistance by blocking the increase in mitochondrial mass and activity induced by ionising irradiation"

**Table S1.** Antibodies used in Western Blot (WB) and immunofluorescence (IF) experiments

| | | Species | Dilution used |
|---|---|---|---|
| Anti-actin | SIGMA (A2066) | Rabbit | 1:1000 |
| Anti-DNA | Progen [AC30-10] | Mouse | 1 :200 |
| Anti-DRP1 | BD bioscience [22/DRP1] | Mouse | 1:1000 |
| Anti-OPA | Cell signalling [D7C1A] | Rabbit | 1:400 |
| Anti-PMPCB / MPPB | Protein tech polyclonal | Rabbit | 1 :250 |
| Anti-TFAM | Cell signalling [D5C8] | Rabbit | 1:1000 |
| Anti-TOM20 | Santa cruz [F10] | Mouse | 1:500 |
| Anti-Vinculine | Abcam Ab18058 | Mouse | 1:4000 (WB) |
| Anti-rabbit Alexa Fluor 647 | Invitrogen, A21245 | Goat | 1:1000 |
| Anti-mouse Alexa Fluor 488 | Invitrogen, A11001 | Goat | 1:1000 |
| Anti-rabbit-Alexa Fluor 594 | Invitrogen A11012 | Goat | 1:1000 |

**Figure S1. Increase in cellular size induced by ionizing irradiation. A)** Flow cytometry analysis of SSC/FSC were analyze from 1 to 4 days after irradiation and illustrate a progressive increase in cell size. **B)** mtDNA copy number quantification by qPCR using two probes against CYTB and ND4 at 48h and 96h after irradiation. Statistical analysis was performed for four independent experiments with one-way ANOVA (***<0,0005). **C)** Mitotracker Green and TMRE were analyzed in small and big cells 72H after irradiation.

**Figure S2. Analysis of mitochondrial morphology. A)** Illustration of the image processing used for the analysis of mitochondrial morphology. **B)** Non irradiated and irradiated cells were fixed and mitochondrial matrix stained with PMPCB, binary images are presented. Scale bar 10 µm**. C)** Quantification of the number and area of mitochondrial braches was performed for 72 non irradiated cells and 91 irradiated cells from four independent experiments.

**Figure S3. Specificity of DRP1 and OPA1 antibodies.** U2OS cells transfected with siRNA targeting DRP1 and OPA1 were used to validate the specificity of the antibodies in **A)** Western blot and **B)** Immunofluorescence experiments. **C)** DRP1 and **D)** OPA1 signals were measured in more than 100 cells from at least two independent experiments. Scale bar 10 µm. Statistical analysis in C and D were performed in GraphPad using Kruskal-Wallis test (***P ≤ 0.001).**E)** Non irradiated and irradiated cells were stained with antibodies against TOMM22 (magenta) and OPA1 (cyan) and Pearson and Mander's correlation coefficients were calculated for 10 cells from 2 independent experiments. Scale bar 10 µm.

**Figure S4. Functional Hyperfusion of mitochondrial network in irradiated cells. A)** movie illustrating the behavior of a cell after microirradiation. **B)** 96H after an irradiation with gamma rays (6 Gy), microirradiation was performed with a 561 nm laser for 50 msec on the region indicated with a dotted square in cells stained with mitotracker green (magenta) and TMRE(cyan). 1 image every 50 sec is shown over a period of 250 sec. Scale bar 10 µm.

**Figure S5. Real time kinetics of RHPS4 subcellular localization in living cells. A)** RHPS4 was added to the culture medium of cells previously stained with Mitotracker deep red, and time lapse was performed for 20 minutes, with an image every 5 min. Fluorescence of RHPS4 was visualized at both the 488 nm and 561 nm channels. **B)** Same cells as in A after fixation with PFA 2%. Scale bar 10 µm.

**Figure S6. Irradiation induced increase in mtDNA. A)** mtDNA was quantified by qPCR using probes against ND4 and CYTB in U2OS non irradiated cells or 96H after a single exposure to X-rays (6 or 10 Gy) in the presence or absence of 0.5 µM RHPS4. Statistical analysis from three independent experiments was performed with one-way ANOVA (****<0,0001). B,C) mtDNA was quantified by qPCR using probes against ND4 and CYTB in T47D and HMLE cells non irradiated or 96H after a single exposure to Gamma-rays (6 Gy) in the presence or absence of 0.5 µM RHPS4. For both a and B data was normalized to the values in non irradiated (NI) cells fixed to 1. Statistical analysis was performed with one-way ANOVA (*<0.05; ***<0,0005). D) Cell sorting results of U2OS cells 96H after a 6 Gy irradiation i the presence or absence of RHPS4 showing the gates used to discriminate small and big cells. E) After sorting, mtDNA was quantified by qPCR using probes against ND4 and CYTB in small and big cells. Statistical analysis was performed with one-way ANOVA (****<0,0001).

**Figure S7. Sequence analysis of mtDNA.** EKLIPSE (Goudenège et al., 2019) was used for analysis and representation of next-generation sequencing (NGS) results of mtDNA for non irradiated cells or 96H after exposure gamma-rays (6 gy) in the presence or absence of RHPS4**.**

**Figure S1**

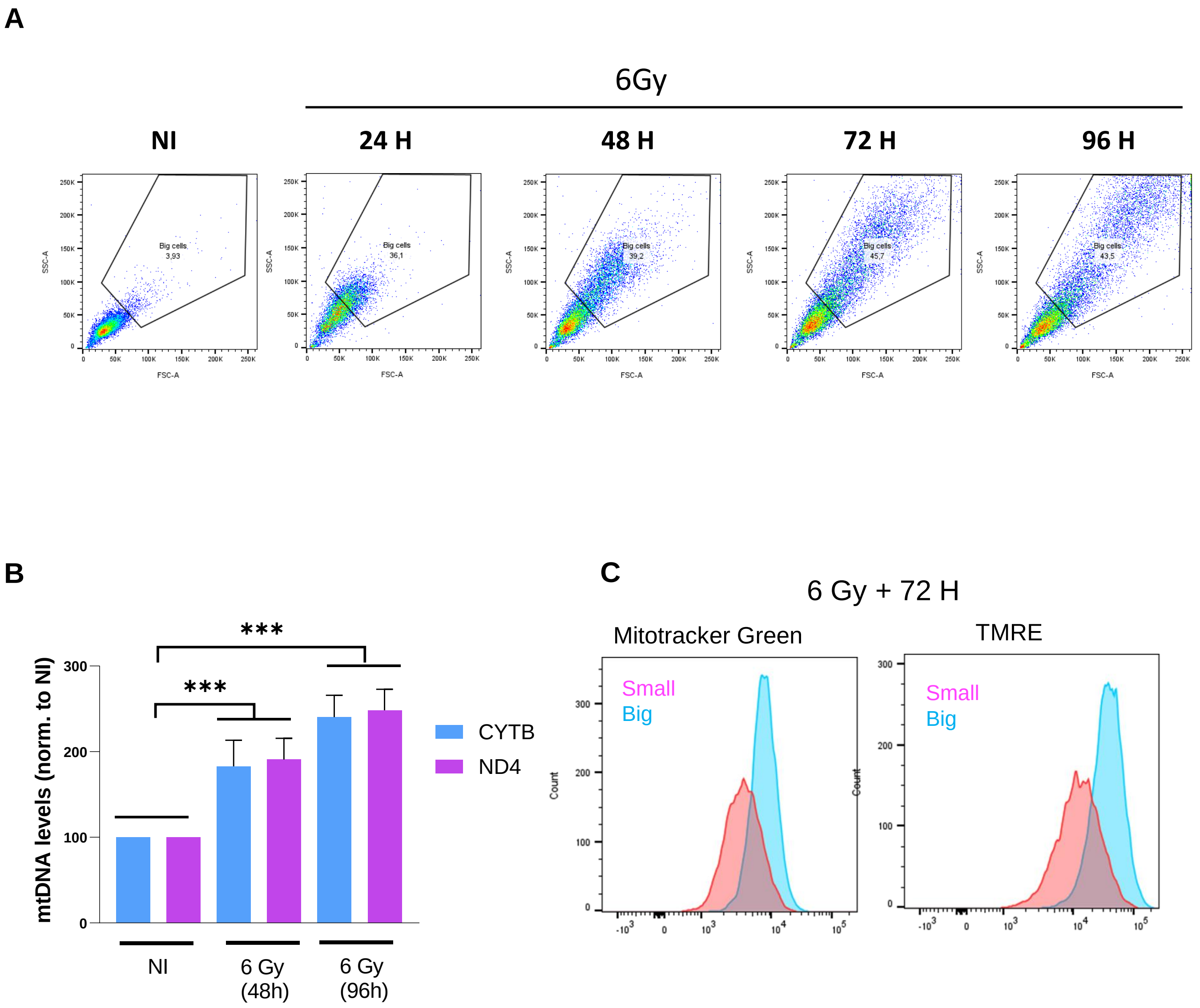
A
6Gy
NI
24 H
48 H
72 H
96 H
Big cells 3,93
Big cells 36,1
Big cells 39,2
Big cells 45,7
Big cells 43,5
SSC-A
FSC-A
B
***
***
mtDNA levels (norm. to NI)
300
200
100
0
CYTB
ND4
NI
6 Gy (48h)
6 Gy (96h)
C
6 Gy + 72 H
Mitotracker Green
TMRE
Small
Big
Count

**Figure S2**

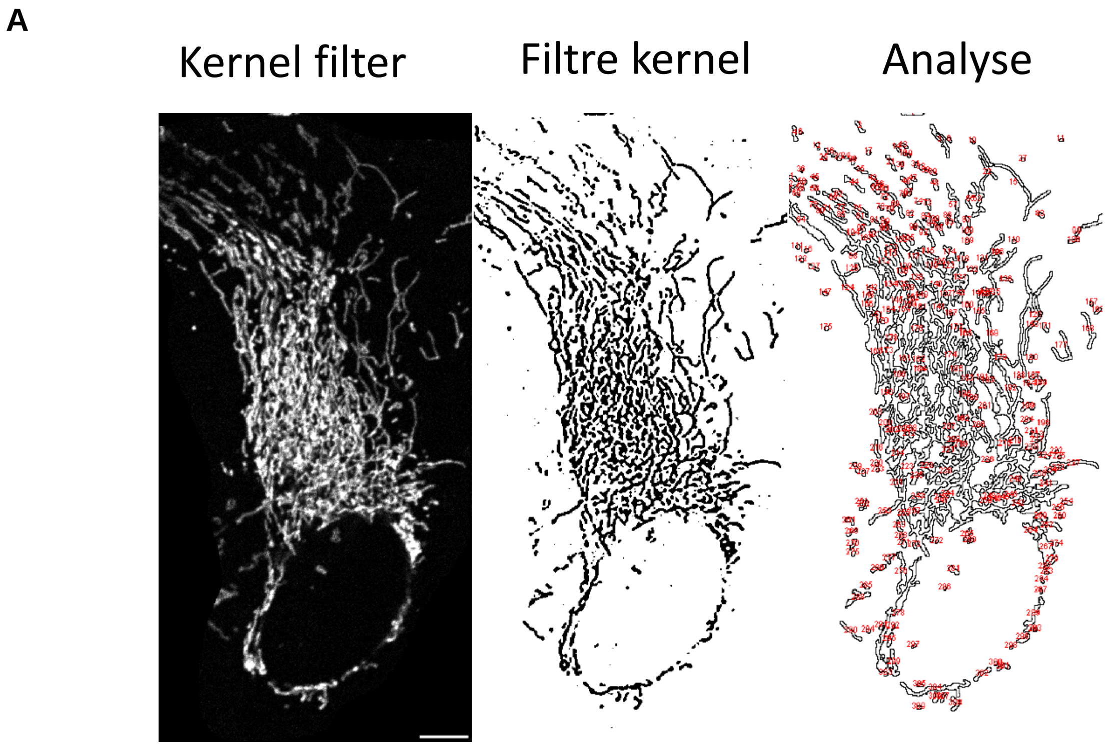
A
Kernel filter
Filtre kernel
Analyse

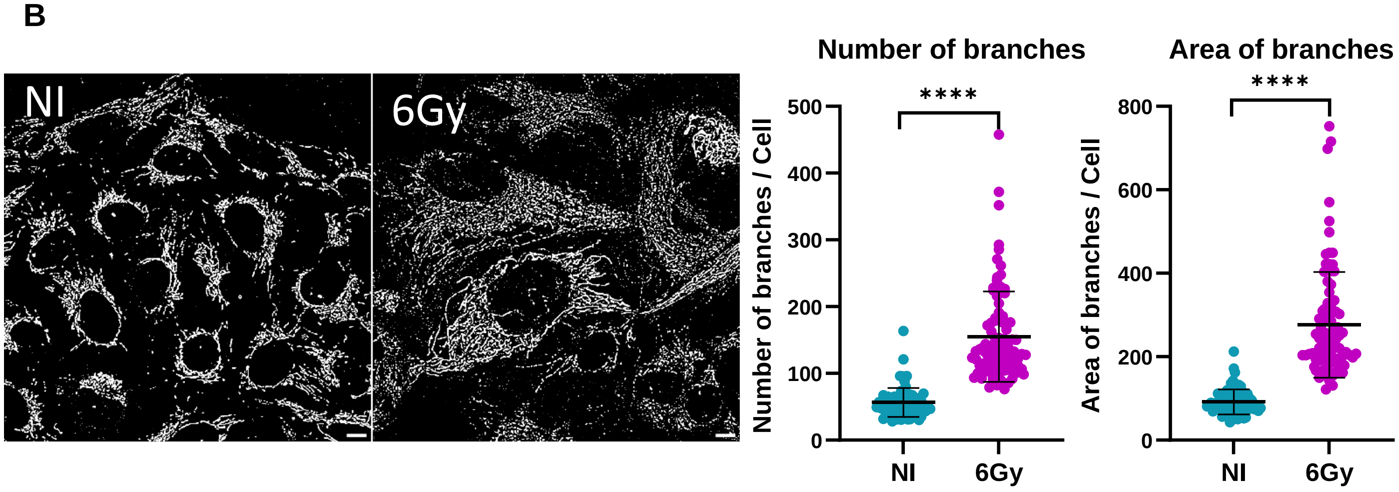
B
NI
6Gy
Number of branches
Number of branches / Cell
****
500
400
300
200
100
0
NI
6Gy
Area of branches
Area of branches / Cell
****
800
600
400
200
0
NI
6Gy

### Figure S3

A

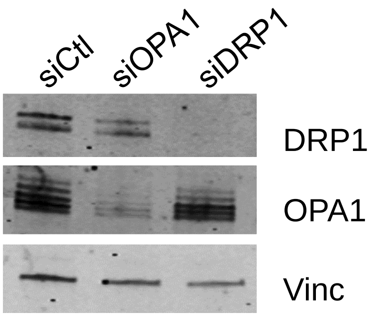


B

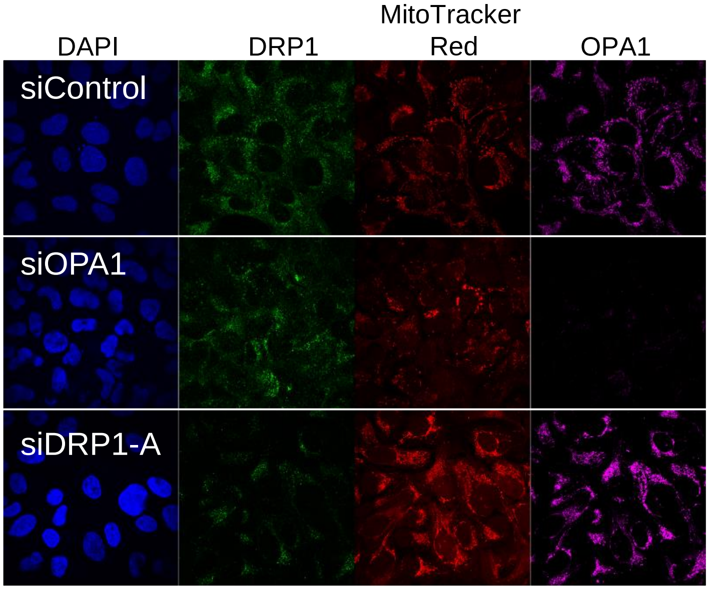


C

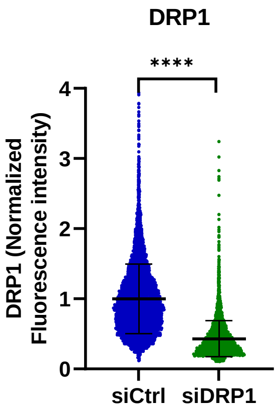


D

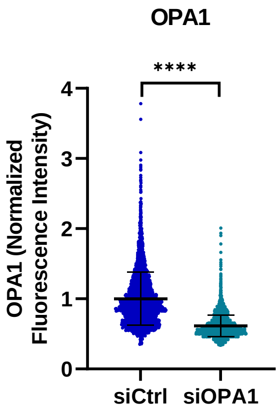


E

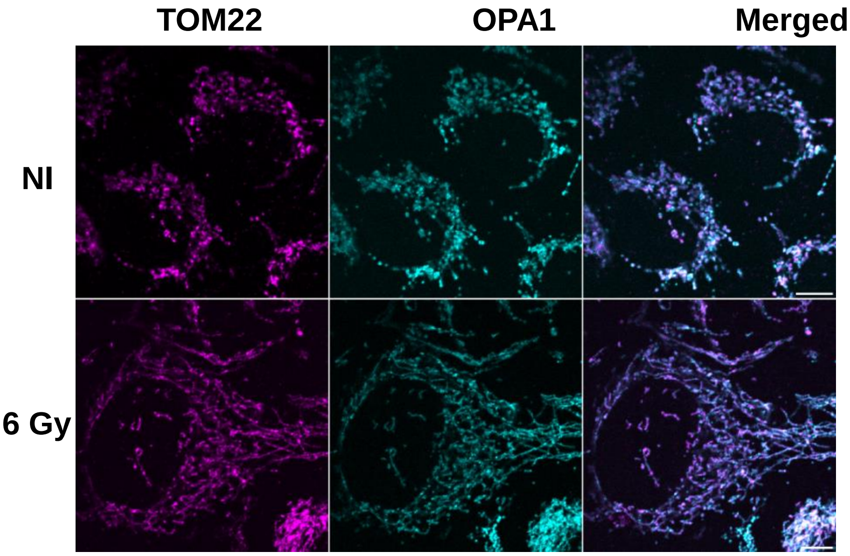


| | Pearson | Mander's |
|---|---|---|
| NI | 0,82 | 0,84 |
| 6 Gy | 0,75 | 0,76 |

**Figure S4**

A

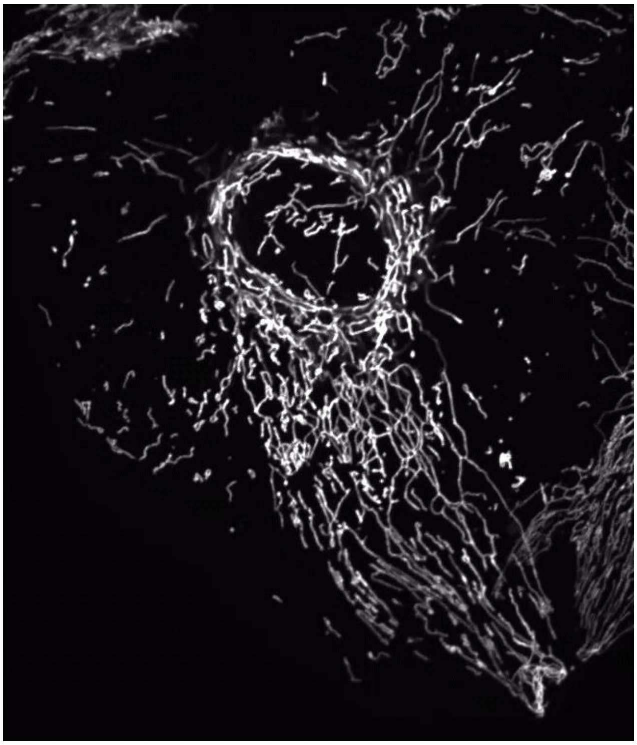

B

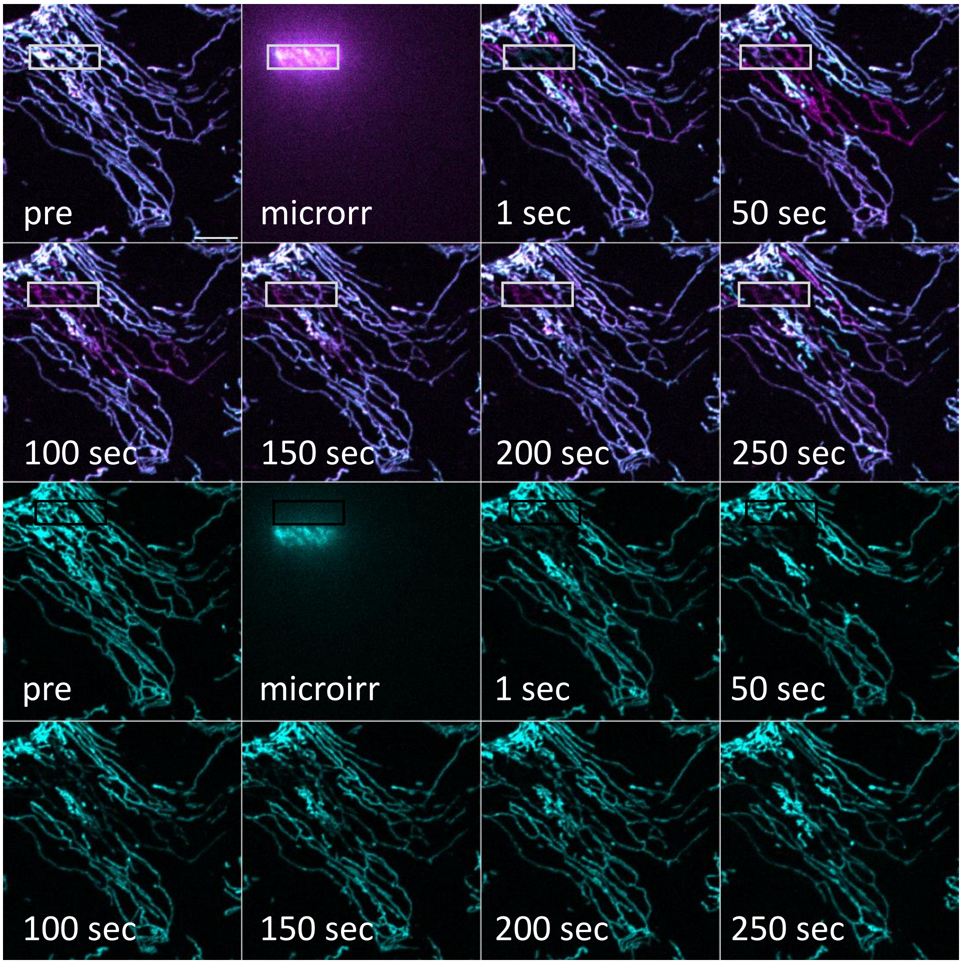
pre
microrr
1 sec
50 sec
100 sec
150 sec
200 sec
250 sec
pre
microirr
1 sec
50 sec
100 sec
150 sec
200 sec
250 sec

**Figure S5**

**A**

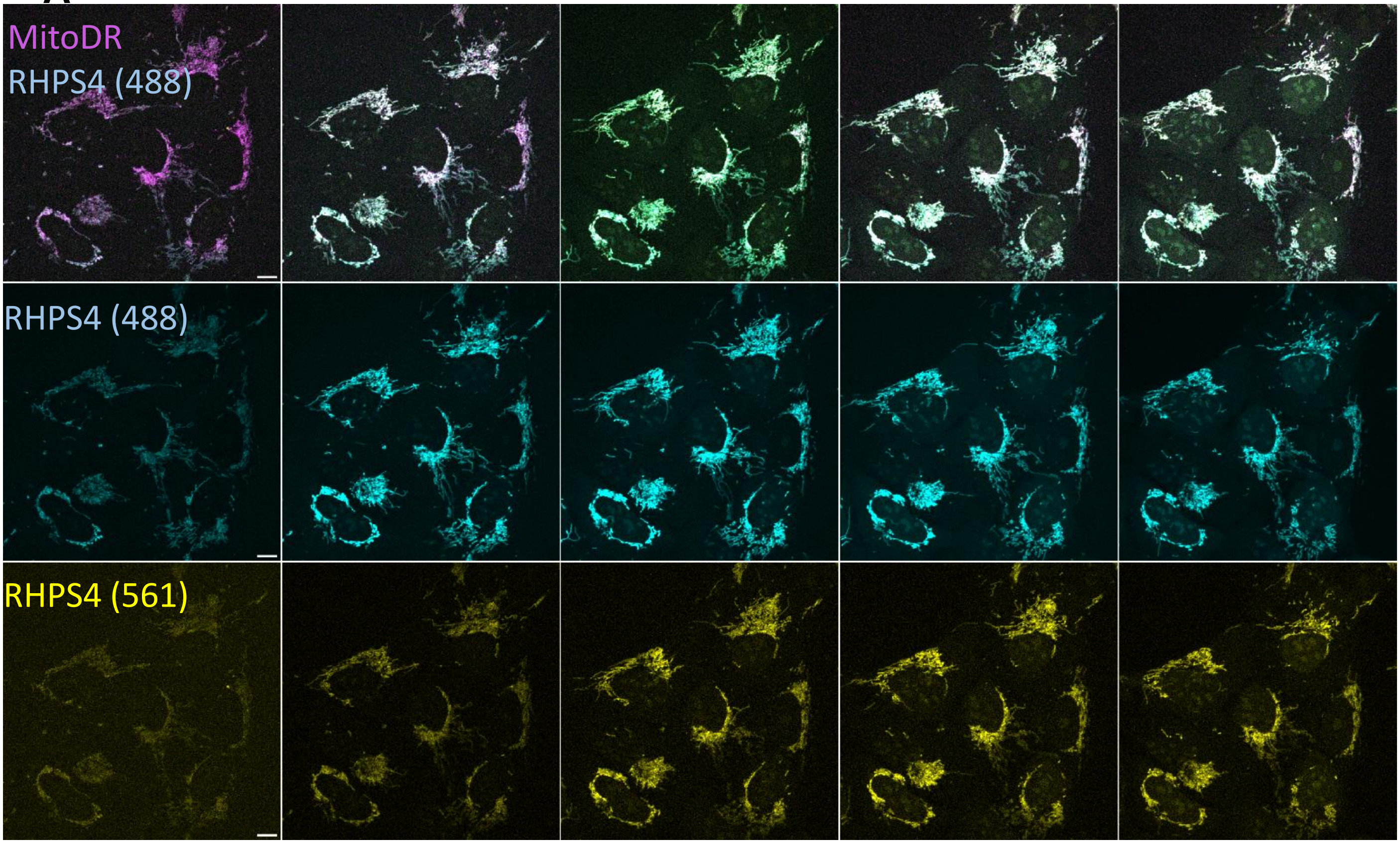


T0 5min 10min 15min 20min

**B**

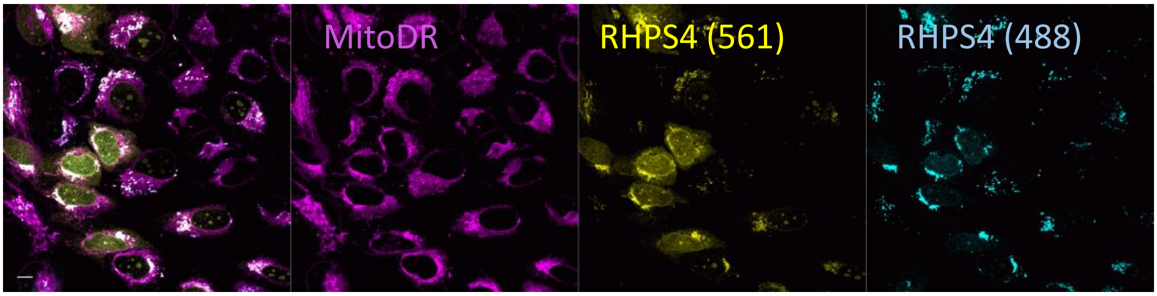

### Figure S6

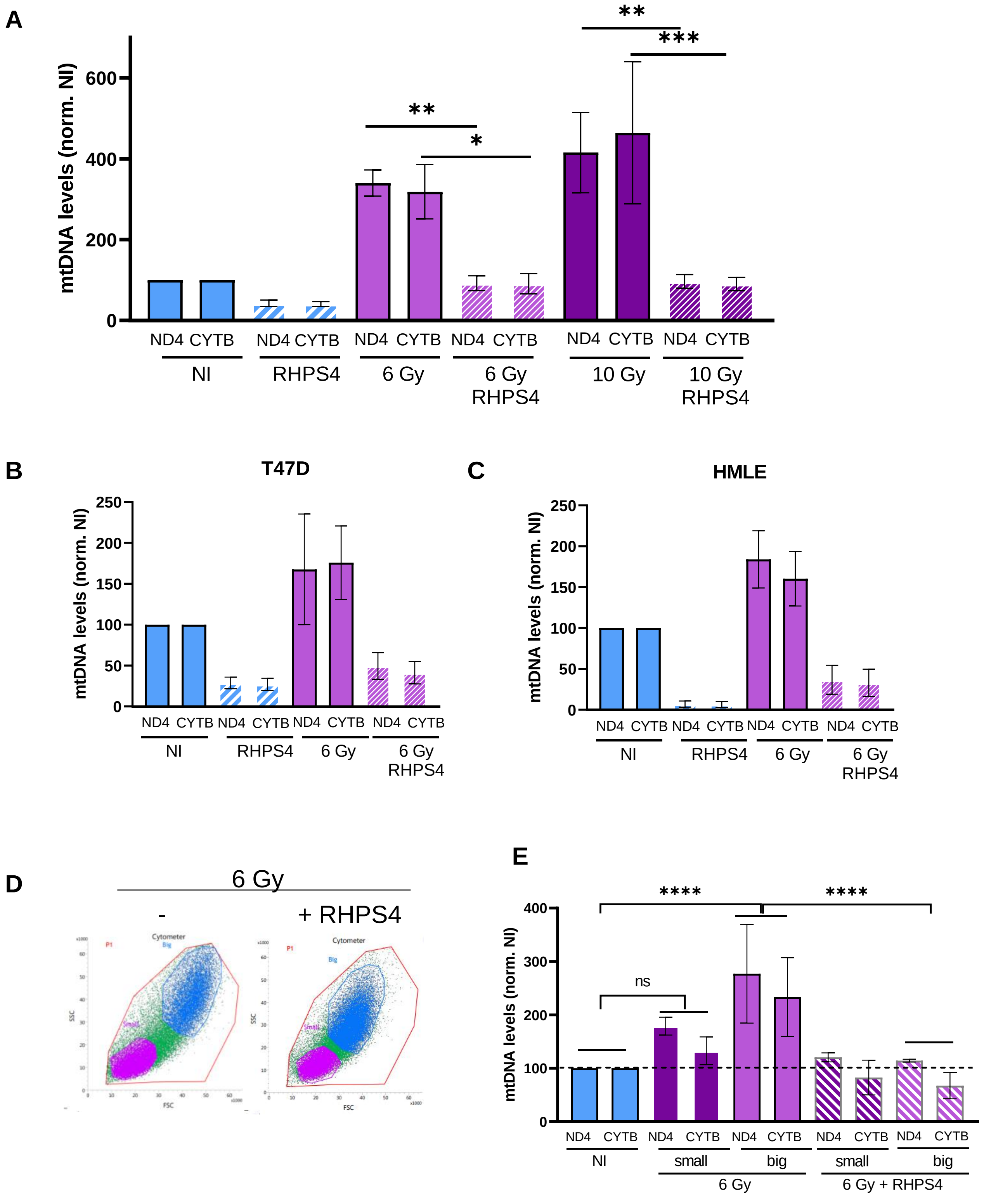

A
mtDNA levels (norm. NI)
0
200
400
600
**
*
**
***
ND4 CYTB ND4 CYTB ND4 CYTB ND4 CYTB ND4 CYTB ND4 CYTB
NI
RHPS4
6 Gy
6 Gy RHPS4
10 Gy
10 Gy RHPS4
B
T47D
mtDNA levels (norm. NI)
0
50
100
150
200
250
ND4 CYTB ND4 CYTB ND4 CYTB ND4 CYTB
NI
RHPS4
6 Gy
6 Gy RHPS4
C
HMLE
mtDNA levels (norm. NI)
0
50
100
150
200
250
ND4 CYTB ND4 CYTB ND4 CYTB ND4 CYTB
NI
RHPS4
6 Gy
6 Gy RHPS4
D
6 Gy
-
+ RHPS4
Cytometer
P1
Big
Small
SSC
FSC
E
mtDNA levels (norm. NI)
0
100
200
300
400
****
****
ns
ND4 CYTB ND4 CYTB ND4 CYTB ND4 CYTB ND4 CYTB
NI
small
big
small
big
6 Gy
6 Gy + RHPS4

### Figure S7

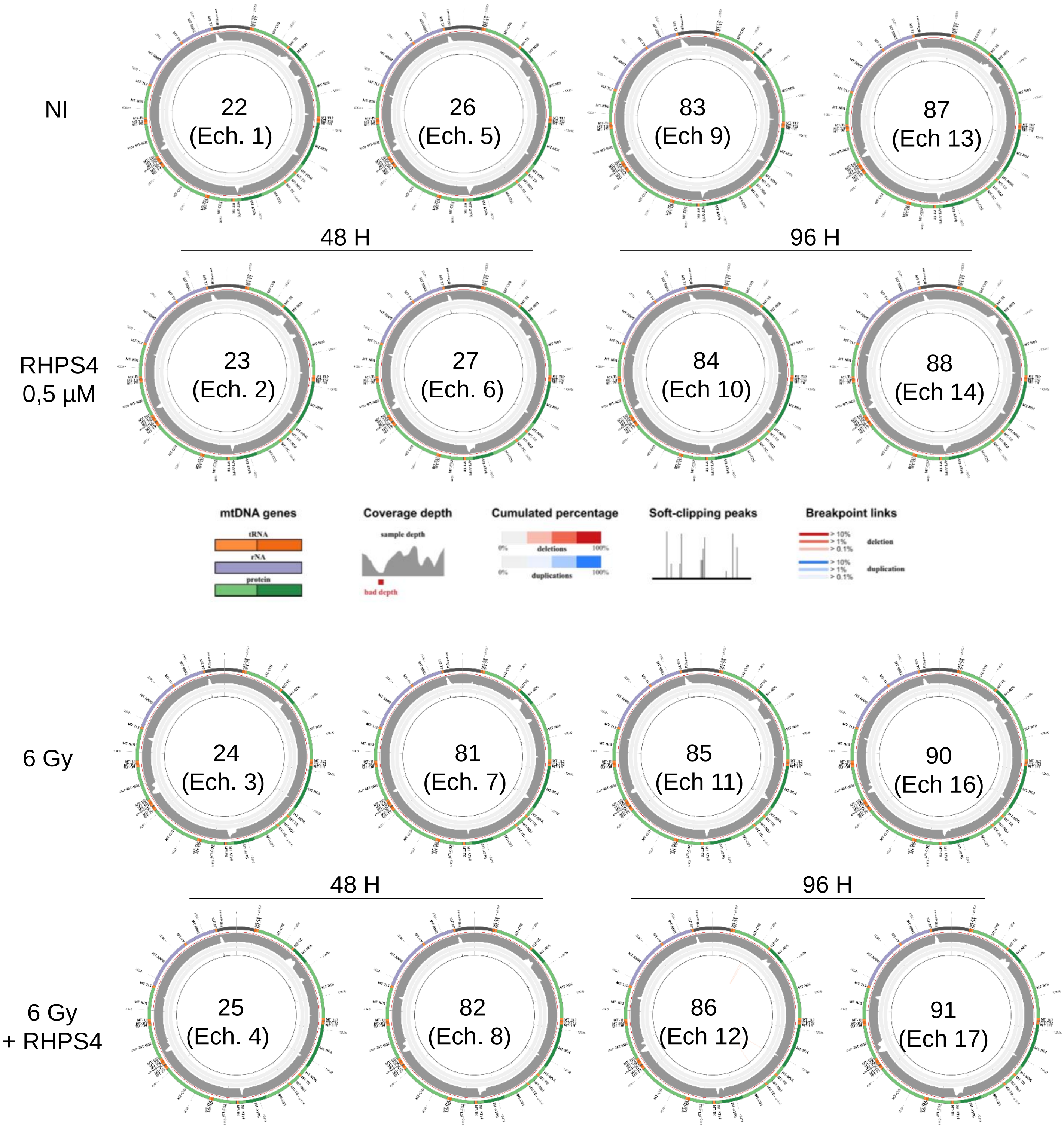
NI
22
(Ech. 1)
26
(Ech. 5)
83
(Ech 9)
87
(Ech 13)
48 H
96 H
RHPS4
0,5 µM
23
(Ech. 2)
27
(Ech. 6)
84
(Ech 10)
88
(Ech 14)
mtDNA genes
tRNA
rNA
protein
Coverage depth
sample depth
bad depth
Cumulated percentage
0%
deletions
100%
0%
duplications
100%
Soft-clipping peaks
Breakpoint links
> 10%
> 1%
> 0.1%
deletion
> 10%
> 1%
> 0.1%
duplication
6 Gy
24
(Ech. 3)
81
(Ech. 7)
85
(Ech 11)
90
(Ech 16)
48 H
96 H
6 Gy
+ RHPS4
25
(Ech. 4)
82
(Ech. 8)
86
(Ech 12)
91
(Ech 17)
